# Crafting for a better MAGIC: systematic design and test for multiparental advanced generation inter-cross population

**DOI:** 10.1101/2021.04.27.441636

**Authors:** Chin Jian Yang, Rodney N. Edmondson, Hans-Peter Piepho, Wayne Powell, Ian Mackay

## Abstract

Multiparental advanced generation inter-cross (MAGIC) populations are valuable crop resources with a wide array of research uses including genetic mapping of complex traits, management of genetic resources and breeding of new varieties. Multiple founders are crossed to create a rich mosaic of highly recombined founder genomes in the MAGIC recombinant inbred lines (RILs). Many variations of MAGIC population designs exist; however, a large proportion of the currently available populations have been created empirically and based on similar designs. In our evaluations of five MAGIC populations, we found that the choice of designs has a large impact on the recombination landscape in the RILs. The most popular design used in many MAGIC populations has been shown to have a bias in recombinant haplotypes and low level of unique recombinant haplotypes, and therefore is not recommended. To address this problem and provide a remedy for the future, we have developed the “magicdesign” R package for creating and testing any MAGIC population design via simulation. A Shiny app version of the package is available as well. Our “magicdesign” package provides a unifying tool and a framework for creativity and innovation in MAGIC population designs. For example, using this package, we demonstrate that MAGIC population designs can be found which are very effective in creating haplotype diversity without the requirement for very large crossing programmes. Further, we show that interspersing cycles of crossing with cycles of selfing is effective in increasing haplotype diversity. These approaches are applicable in species which are hard to cross or in which resources are limited.

## Introduction

The multiparental advanced generation inter-cross (MAGIC) population was initially proposed in crops by Mackay and Powell (2007) as a highly recombined population derived from multiple founders. The first MAGIC population was produced using 19 founders in *Arabidopsis thaliana* (Kover et al. 2009). The MAGIC pedigree described by Cavanagh et al. (2008) has served as a foundation for the design of many MAGIC populations in subsequent years. Briefly, the MAGIC pedigree shows a single funnel going from 8 founders to a recombinant inbred line (RIL). Starting with 8 founders labelled as A to H, two-way crosses are made as (A × B), (C × D), (E × F) and (G × H). Next, four-way crosses are made as ((A × B) × (C × D)) and ((E × F) × (G × H)). Lastly, eight-way crosses are made as (((A × B) × (C × D)) × ((E × F) × (G × H))) followed by several generations of selfing. Using this crossing scheme, the end of the funnel is a RIL with its genome composed of contributions from all 8 founders. Alternatively, a MAGIC population design may involve multiple funnels like the elite wheat MAGIC population by Mackay et al. (2014). Regardless of the designs, MAGIC RILs have diverse recombination landscape and rich mosaics of founder genomes (Scott et al. 2020).

Over the years, MAGIC populations have been used in various studies with great success. MAGIC populations are popular choices in mapping quantitative trait loci (QTLs), for examples, resistance QTLs in bread wheat (Stadlmeier et al. 2019), cold tolerance QTLs in maize (Yi et al. 2020) and high-throughput phenotype QTLs in rice (Ogawa et al. 2021). In addition to single trait analyses, multivariate analyses (multi-trait or multi-environment) have been demonstrated in MAGIC populations (Scutari et al. 2014, Verbyla et al. 2014). Diouf et al. (2020) used a tomato MAGIC population to dissect the underlying genetic-by-environment (G × E) and plasticity for climate adaptation traits. Aside from QTL mapping, MAGIC populations are valuable resources for genomic selection owing to their properties of highly recombined genomes and large population size (Scott et al. 2021). Following that, there are opportunities for using MAGIC RILs in breeding new varieties (Bandillo et al. 2013, Li et al. 2013). With large numbers of founders, MAGIC populations also provide a dynamic asset for the management of genetic resources (Thépot et al. 2015) and may be used to improve our understanding of crop adaptation (Scott et al. 2021). Given the longevity with a broad array of uses, MAGIC populations are an invaluable community resource for creative and impactful research.

Considering the importance of MAGIC populations in crop research, we sought to understand the relationship between MAGIC population designs and population recombination landscape. We selected and analyzed five MAGIC populations with publicly available marker genotypes for the founders and RILs, genetic map positions and pedigrees. The selected populations comprise the UK wheat 8-founder (Mackay et al. 2014), German wheat 8-founder (Sannemann et al. 2018), cowpea 8-founder (Huynh et al. 2018), tomato 8-founder (Pascual et al. 2015) and UK wheat 16-founder (Scott et al. 2021) MAGIC populations. These MAGIC populations were created from different designs. We contrasted the observed recombinant haplotypes to expected (simulated) recombinant haplotypes in each population. A comparable cross-population analysis can be challenging due to many variables like genome sizes, marker genotyping platforms and numbers of founders. Fortunately, there are two elite wheat 8-founder MAGIC populations (Mackay et al. 2014, Sannemann et al. 2018) that were genotyped with the same 90k SNP array (Wang et al. 2015), which allowed us to directly compare the two populations in greater depth. We found that the recombination landscape varies across designs and that this variation is consistent across species.

Following our results, we have created the “magicdesign” package in R (R Core Team 2021) for the purpose of creating and testing different MAGIC population designs. Three major steps are involved in the package pipeline: design creation, population simulation and comparative analysis. Users can create a design by either specifying input variables or providing a custom pedigree. Once a design is created, “magicdesign” converts it into a crossing scheme from the founders to final RILs and simulates a population based on the crossing scheme. After multiple iterations of simulation, distributions of recombinant haplotypes and founder genomes are summarized. Results from one or more designs can be combined and compared visually in plots. In addition, “magicdesign” produces a pedigree in both text and plot formats that can be used as a guide to support crossing work in practice. Aside from the described roles, “magicdesign” serves as a tool to advance the use of MAGIC in future multiparental populations.

## Materials and Methods

### Evaluation of MAGIC population designs

We surveyed all available MAGIC populations that have been published to date (including pre-prints) and identified five populations with publicly available marker data. These five MAGIC populations include wheat with 8 UK elite founders (Mackay et al. 2014), wheat with 8 German elite founders (Sannemann et al. 2018), cowpea with 8 founders (Huynh et al. 2018), tomato with 8 founders (Pascual et al. 2015) and wheat with 16 UK diverse founders (Scott et al. 2021). These populations are referred to as wheat-UK8, wheat-DE8, cowpea, tomato and wheat-UK16, respectively (Table 1). These datasets were chosen because the marker data for the founders and recombinant inbred lines (RILs), genetic map positions, and pedigree are publicly available. The wheat-UK16 population is an exception as it has founder genotype probabilities to compensate for the lack of genetic map positions. Links to the source datasets are listed in Supplementary Text. Other publicly available datasets like the wheat with 8 German founders (Stadlmeier et al. 2018), wheat with 8 Australian founders (Shah et al. 2019), rice with 8 founders (Raghavan et al. 2017), maize with 9 founders (Dell’Acqua et al. 2015) and *Arabidopsis* with 19 founders (Kover et al. 2009) are excluded because at least one component of the data needed for our purpose is not present.

**Table 1.** Summary of five analyzed MAGIC populations. The wheat-UK8 and wheat-DE8 datasets have been reduced to share the same markers and maps for comparison. The proportion of recombinant haplotypes recovered (PRHR) is calculated as number of recombinant haplotypes in actual dataset divided by number of recombinant haplotypes in simulated dataset. PRHR is shown as mean ± standard deviation.

| dataset | n | genome<br>(cM) | marker | distance<br>(cM/marker) | PRHR | reference |
| --- | --- | --- | --- | --- | --- | --- |
| wheat-UK8 | 8 | 5,262 | 5,138 | 1.024 | 0.296 ± 0.094 | Mackay et al. (2014) |
| wheat-DE8 | 8 | 5,262 | 5,138 | 1.024 | 0.331 ± 0.058 | Sannemann et al. (2018) |
| cowpea | 8 | 979 | 32,114 | 0.030 | 0.674 ± 0.207 | Huynh et al. (2018) |
| tomato | 8 | 2,156 | 1,345 | 1.603 | 0.410 ± 0.106 | Pascual et al. (2015) |
| wheat-UK16 | 16 | 5,262 | 1,065,178 | 0.005 | 0.799 ± 0.092 | Scott et al. (2021) |

All five chosen MAGIC populations vary in numbers of RILs and marker density (Table 1). The original wheat-UK8 dataset is made of 643 RILs and 18,599 markers while the original wheat-DE8 dataset is made of 910 RILs and 7,579 markers. To maintain a fair comparison between these two populations, we kept only 5,138 markers that are common between wheat-UK8 and wheat-DE8. In wheat-DE8, missing data were previously imputed to numerical mean (twice the allele frequency). These imputed marker data cannot be used in “qtl2” (Broman et al. 2019) for calculating founder genotype probabilities, so we reverted the imputed marker data by converting any non-integer marker data to missing. The cowpea dataset is made of 305 RILs and 32,114 markers after removing 16 markers in the original dataset where the marker data is missing in at least one founder. The tomato dataset is made of 238 RILs and 1,345 markers. The wheat-UK16 dataset is made of 504 RILs and 1,065,178 markers.

### Identification of recombinant haplotypes in MAGIC populations

To identify recombinant haplotypes, the biallelic marker data in the RILs need to be converted into founder genotypes. For each dataset except wheat-UK16, we determined the founder genotypes in each RIL using the “qtl2” package (Broman et al. 2019) in R (R Core Team 2021). We first calculated the founder genotype probabilities using *calc.genoprob* function with error probability of 0.01 (1%) and Haldane map function. Next, we inferred the founder genotypes from the probabilities using *maxmarg* function with minimum probability of 0.5001. We chose a slightly higher threshold than the previously used minimum probability of 0.5 by Gardner et al. (2016). Since the genotype probabilities for all founders at each RIL’s marker sum to 1, the threshold of 0.5001 eliminates the risk of the *maxmarg* function picking a founder genotype at random when there are two or more with the same probability above the threshold. For the wheat-UK16 dataset, the founder genotype probabilities are readily available. These probabilities were calculated from STITCH (Davies et al. 2016), which is a different software but uses the same underlying hidden Markov model (HMM) as “qtl2”. We inferred the founder genotypes in the wheat-UK16 dataset using an equivalent threshold of 1 because the probabilities are provided as a sum of two allele probabilities. Markers without any founder genotype probabilities above the threshold were set to missing.

Using the inferred founder genotypes in each dataset, we identified the recombinant haplotypes at each breakpoint. The recombinant haplotype is a combination of flanking founder genotypes at each breakpoint. For example, 1_5 is the recombinant haplotype at a breakpoint where the two flanking founder genotypes are founder 1 and 5. For a population with *n* founders, there are *n*^2^ − *n* individual recombinant haplotypes. Therefore, in an 8-founder MAGIC population, there are 56 individual recombinant haplotypes (1_2, 1_3, …, 8_6, 8_7). In order to summarize the results, we summed the counts of each individual recombinant haplotype in every RIL, and averaging the counts across all RILs to obtain the mean counts of individual recombinant haplotypes.

As a control, we simulated a similar MAGIC population based on the original pedigree for each dataset and calculated the true counts of recombinant haplotypes. We first derived an approximated crossing scheme from the pedigree. Since the information on replicates is not always present in the pedigree, we assumed that no replicates and considered all funnels to be independent. Next, we used the “AlphaSimR” package (Gaynor et al. 2021) in R (R Core Team 2021) to simulate the MAGIC populations for a total of 100 iterations. In order to keep track of founder genotypes, we expanded each marker into *n* − 1 markers with the same exact genetic map position to prevent recombination among these markers. Using *n* = 4 founders as example, the founders are coded as 000, 100, 010 and 001 across the three expanded markers. This expanded marker system tracked the true founder genotypes from the start to the end of simulation, and therefore allowed us to calculate the true counts of recombinant haplotypes using the same method described in the previous paragraph. In all datasets except for wheat-UK16, we used the same genetic map positions as the actual datasets. In wheat-UK16, we used equally-spaced markers at 0.5 cM since the genetic map positions were not available for this dataset.

We included an additional control using a hybrid approach of actual and simulated datasets. Specifically, we converted the founder genotypes in simulated RILs into biallelic marker data and inferred the founder genotypes using the same procedures as we did in the actual datasets. This approach was applied to all four MAGIC datasets except wheat-UK16. The counts of recombinant haplotypes identified from this approach provide an upper limit to the inferred number of founder genotypes using “qtl2” (Broman et al. 2019). There is a caveat, however, that since our simulation is based on an approximated crossing scheme, the outcomes of this approach may not precisely represent the upper limit.

### Determination of unique or identical recombinant haplotypes

Following the identified recombinant haplotypes in wheat-UK8 and wheat-DE8, we classified these recombinant haplotypes into unique or identical groups based on their positional overlaps within an interval. Recombinant haplotypes of the same founder pairs are considered identical if they fall within the same interval, otherwise unique if they do not overlap. The intervals are arranged in non-overlapping bins of approximately 1 or 10 cM from the start to end of chromosome. Identical recombinant haplotypes exist due to replications of cross progeny in the MAGIC population. While this classification cannot distinguish between identical and independent recombinant haplotypes within the interval, the probability of independent recombinant haplotypes is low and assumed equal between wheat-UK8 and wheat-DE8. Therefore, the results from this comparison can elucidate the effect of MAGIC population designs on the proportions of unique against identical recombinant haplotypes.

### Minimum probability in calling founder genotypes

The minimum probability used in calling founder genotypes determines the power in identifying the correct founder genotypes in the RILs. We selected 10 thresholds ranging from 0.1 to 1.0 with an increment of 0.1. We applied each threshold to the *maxmarg* function in “qtl2” (Broman et al. 2019) in simulated populations based on wheat-UK8 and wheat-DE8. These simulated populations are similar to the previously described hybrid approach where the simulated founder genotypes in RILs are converted to biallelic markers prior to calculating genotype probabilities. For each threshold, we computed the proportions of correct, incorrect or missing founder genotypes by comparing the inferred to true founder genotypes. The ideal threshold should have a high proportion of correct founder genotypes with low proportions of incorrect and missing founder genotypes.

### Marker density in MAGIC population

We used the proportion of recombinant haplotype recovered (PRHR) to quantify the recombinations in MAGIC population that is captured in the marker data. We calculated the empirical PRHR in all five datasets as the counts of recombinant haplotypes in the actual dataset divided by the counts in the simulated dataset. Since the marker density is not constant along the genomes in actual datasets, we sought to determine a clearer relationship between PRHR and marker density via simulation. We simulated a single chromosome of 200 cM with 8 founders crossed using the same design as wheat-UK8. The founder alleles were simulated based on the correlations among founders in wheat-UK8 using the *rmvbin* function in “bindata” package (Leisch et al. 2021) in R (R Core Team 2021). We simulated a total of 4,000 markers that are equally spaced at 0.05 cM. To test lower marker densities, we thinned the same simulated marker data to 0.10, 0.20, 0.40, 0.80, 1.60, 3.20, 6.40, 12.80 cM respectively. For each marker density, we inferred the founder genotypes in the RILs using “qtl2” (Broman et al. 2019) at a probability threshold of 0.5001 and calculated the counts of recombinant haplotypes. Lastly, we obtained the PRHR by taking the counts of recombinant haplotypes for each marker density divided by the true counts.

### Data Availability

The “magicdesign” package and its installation instructions are available for download at https://github.com/cjyang-sruc/magicdesign. Detailed instructions are available at https://cjyang-sruc.github.io/magicdesign_vignette. The Shiny app “magicdesignee” can be found at https://magicdesign.shinyapps.io/magicdesignee/. R scripts used in all analyses can be found at https://cjyang-sruc.github.io/files/magicdesign_analysis.R.

## Results

### Classifications of MAGIC population designs

Variations in the MAGIC population designs can be described by the number of founders and the crossing scheme (Figure 1A). It is convenient to first consider two classes of designs based on the number of founders: power of two (P2) and non-power of two (NP2). As the names suggest, the P2 class has *n* = 2^*i*^ founders for any *i* > 1 whereas the NP2 class has *n* ≠ 2^*i*^ and *n* > 2 founders. P2 designs are generally easier to implement in practice because the numbers of individuals in a funnel are halved in every crossing generation. For either P2 or NP2 classes, the crossing scheme can be structured, unstructured and semi-structured (Figure 1A). A structured design involves strictly defined crosses among the founders and intermediaries such that the crossing scheme can be further classified into full, partial balanced, partial unbalanced or basic designs. These designs are elaborated further in subsequent paragraphs. On the other hand, an unstructured design involves random crosses among the founders and intermediaries, while a semi-structured design is a combination of structured and unstructured designs. Additional features of a structured design include: (1) precise tracing of the ancestry of each RIL back to its progenitors, (2) number of crossing generations is equal to *log*_2_*n* rounded up to the nearest integer. These features may not hold true in unstructured or semi-structured designs.

**Figure 1.**
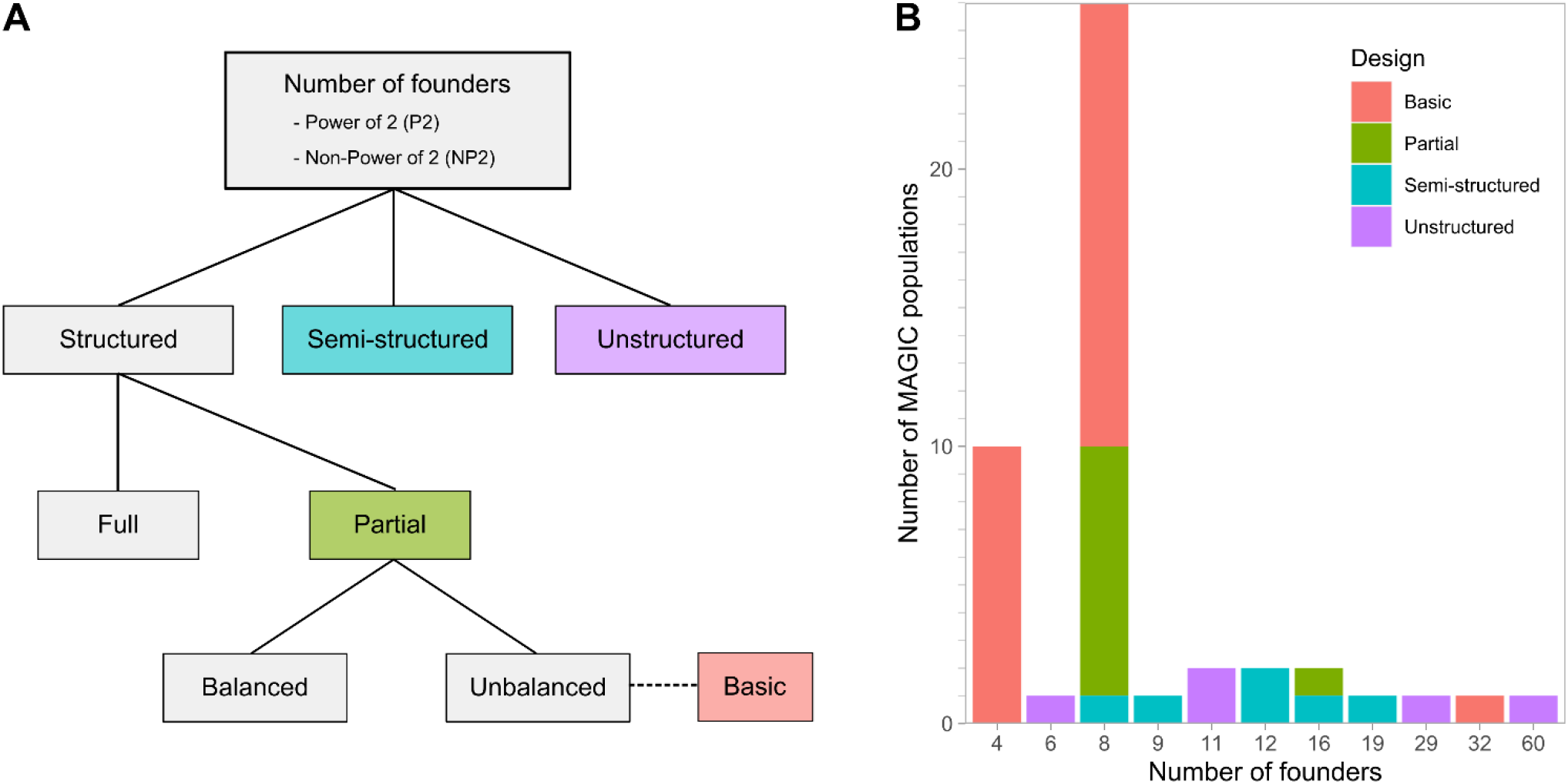
Classifications of MAGIC population designs. [**A**] Flowchart of classifying MAGIC population designs based on their crossing schemes. [**B**] Distribution of MAGIC population designs in all 48 surveyed populations.

Within a structured design, there are two primary types based on the number of funnels: full and partial (Figure 1A). A full P2 design has *n*!⁄2^*n*−1^ funnels while a partial P2 design has one or more funnels but less than *n*!⁄2^*n*−1^ funnels. The numerator is the total number of permutations of 1 to *n* founders, and the denominator is the total number of equivalent permutations by the MAGIC definition. Directions of crosses are disregarded in defining a funnel. The denominator can be described as 2^*n*−1^ = ∏ 2^*n*⁄*x*^ where *x* = 2^*i*^ ≤ *n* for *i* > 0. In an example with four founders, 1234, 1243, 2134, 2143, 3412, 3421, 4312 and 4321 are all equivalent funnels. For simplicity, ((1 × 2) × (3 × 4)) is written as 1234. Full and partial types exist in NP2 designs although the number of funnels in a full design cannot be generalized similarly (Table S1). From a practical perspective, a full P2 design is achievable for four or eight founders but not for 16 or more founders as the number of required funnels becomes unmanageable.

Within a partial design, the funnels can be chosen in either a balanced or an unbalanced way (Figure 1A). A balanced design has an equal number of founders among the funnels and equal frequency of founder pairs at each crossing generation. In a four-founder MAGIC design, 1234, 1324 and 1423 form a set of three balanced funnels. First, each founder occurs thrice in the set of funnels. Second, each founder is paired once with another founder in the two-way crosses, and twice in the four-way crosses. For example, founder 1 meets founder 2 once in the two-way cross (first funnel) and twice in the four-way cross (second and third funnels). Coincidentally, since *n* = 4 and *n*!⁄2^*n*−1^ = 4!⁄2^3^ = 3, the set of three balanced funnels is equivalent to a full design for four founders. Unlike the partial balanced design where the number of funnels is restricted to set rules, the partial unbalanced design is formed by funnels chosen randomly. Differences between balanced and unbalanced designs are explored in a later section. Additionally, we coin the special case of partial unbalanced design with one funnel as a basic design. Examples of all of the designs are shown in Figure S1.

Based on our survey of 48 MAGIC populations in 15 crop species that have been described in either published or pre-print literature to date, there are 39 P2 and 9 NP2 designs (Figure 1B and Table S2). The numbers of founders range from four to 60, with 4 and 8 founders as the predominant numbers. Despite the ease of handling required for crosses based on a full design with 4 founders, all 10 of the populations were created using a basic design. Of the 26 MAGIC populations with 8 founders, there are 16 basic designs, nine partial designs and one semi-structured design. The popularity of the basic design can be ascribed to Cavanagh et al. (2008), who provided an illustrated pedigree of a basic design. We refrained from classifying the partial designs into balanced and unbalanced designs due to the lack of pedigree information in many MAGIC populations. Regardless of the number of founders, there has not been any MAGIC population created with a full design. There are several 8-founder populations that came close to a full design. The bread wheat MAGIC population by Mackay et al. (2014) had 210 out of 315 required funnels for a full design. The maize MAGIC population by Dell’Acqua et al. (2015) had mixed funnels from pooling different four-way individuals and had to introduce an additional founder due to a failed two-way cross. The three bread wheat MAGIC populations by Shah et al. (2019) came closest to a full design with 311 to 313 funnels.

**Table 2.**
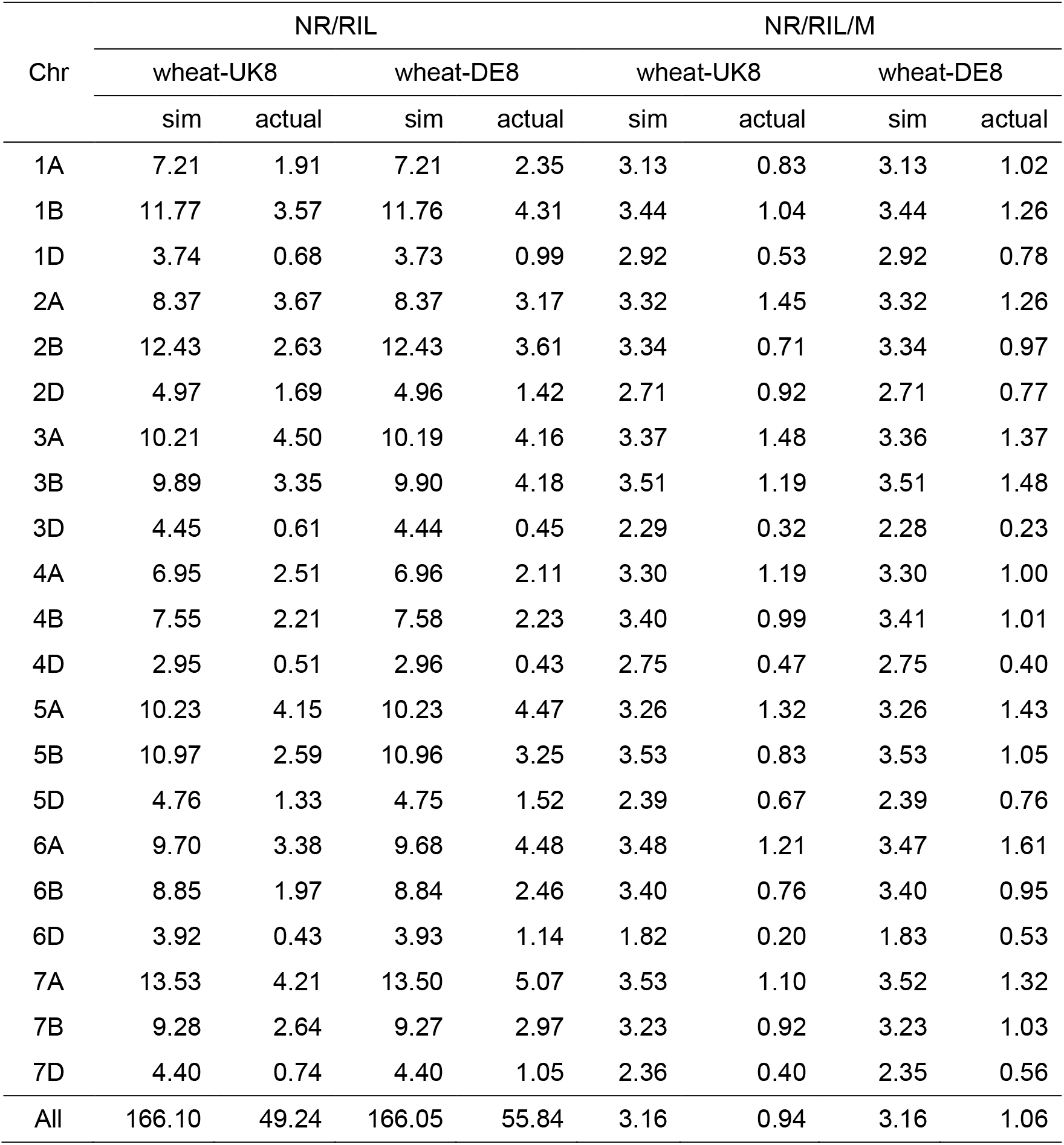
Number of informative recombinations in wheat-UK8 and wheat-DE8. The number of informative recombinations (NR) is calculated for both simulated and actual wheat-UK8 and wheat-DE8 datasets. Note: recombinant inbred line (RIL), Morgan (M).

### Empirical evaluation of two bread wheat MAGIC populations

Our evaluation on two bread wheat MAGIC populations derived from distinct sets of 8 elite founders shows that the distributions of recombinant haplotypes differ for each MAGIC design (Figure 2 and Figure S2). We used the wheat-UK8 and wheat-DE8 populations, in which wheat-UK8 is an example of a partial design while wheat-DE8 is an example of a basic design (Table S2). To maintain our cross-population comparison as fair as possible, we reduced the original wheat-UK8 and wheat-DE8 datasets to smaller subsets with common markers (Figure 2), although the same analysis was performed on the original datasets too (Figure S2). The subsets include all 643 RILs in wheat-UK8 and 910 RILs in wheat-DE8, and 5,138 common markers arranged in the same genetic map positions as Gardner et al. (2016). This genetic map is chosen over the original genetic map in the wheat 90k array (Wang et al. 2014) because of higher map quality.

**Figure 2.**
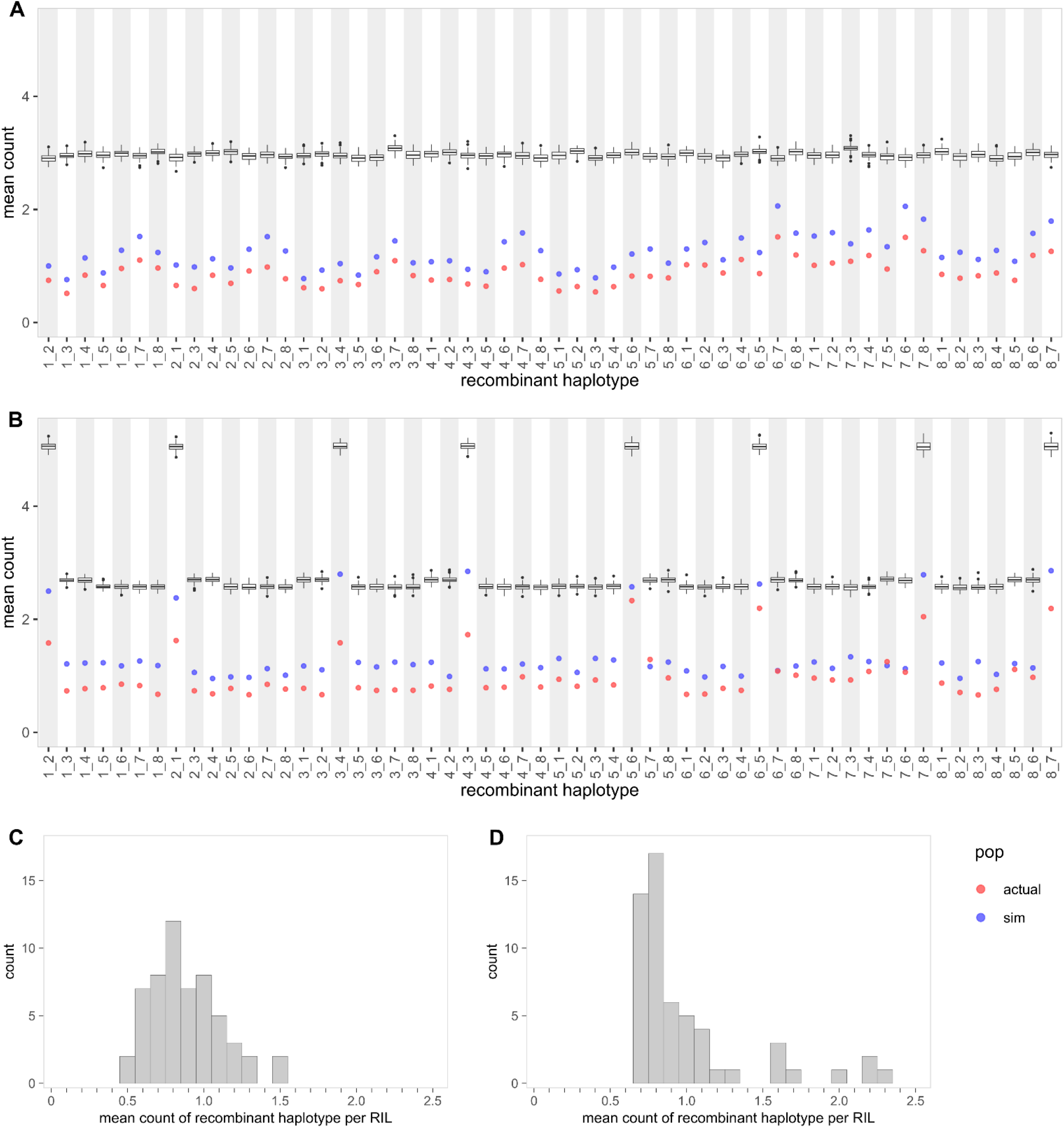
Distributions of recombinant haplotypes in two wheat MAGIC populations. [**A**] Plot shows mean count of each recombinant haplotype in a single RIL in wheat-UK8. The boxplot shows mean count from true founder genotypes (100 simulated iterations). The red and blue points show mean count from inferred founder genotypes. [**B**] Plot shows mean count of each recombinant haplotype in a single RIL in wheat-DE8. [**C**] Histogram of the mean count in wheat-UK8. [**D**] Histogram of the mean count in wheat-DE8.

The distribution of all recombinant haplotypes is less skewed in wheat-UK8 than in wheat-DE8 (Figure 2). In wheat-UK8, none of the recombinant haplotype appears more frequently than others (Figure 2A). In any given RIL, there are 0.879 ± 0.227 (mean ± standard deviation) individual recombinant haplotypes. In wheat-DE8, eight recombinant haplotypes appear about twice as frequently as the others (Figure 2B). There are 1.910 ± 0.313 of these eight recombinant haplotypes (1_2, 2_1, 3_4, 4_3, 5_6, 6_5, 7_8 and 8_7) instead of 0.845 ± 0.150 of the other recombinant haplotypes. In addition, the mean count of recombinant haplotype is approximately normally distributed in wheat-UK8 (Figure 2C) but is skewed to the right in wheat-DE8 (Figure 2D). The eight skewed recombinant haplotypes match with all of the founder pairs in two-way crosses in wheat-DE8. This is not a coincidence because two-way crosses have the largest founder genomes to recombine. With every generation of crosses, the founder genomes are halved and so there are less recombinations between any two founders.

While wheat-UK8 has a slightly lower number of recombinant haplotypes per RIL than wheat-DE8 in both reduced (Table 2) and full (Table S3 and S4) datasets, the proportion of unique recombinant haplotypes is higher in wheat-UK8 than in wheat-DE8 (Figure 3 and Figure S3). Due to the imprecision of inferred recombination breakpoints, we defined recombinant haplotypes with breakpoints within any non-overlapping intervals as identical. We chose the intervals to be 1 cM and 10 cM wide. With the interval width set to 1 cM, the number of identical recombinant haplotypes in wheat-UK8 ranges from 1 (unique) to 21 (Figure 3A) while the number in wheat-DE8 ranges from 1 (unique) to 128 (Figure 3B). In addition, 11,602 out of 31,642 recombinant haplotypes in wheat-UK8 are considered unique, and 9,567 out of 50,794 recombinant haplotypes in wheat-DE8 are considered unique (Figure 3C). After discounting the identical recombinant haplotypes, there are 17,786 distinct recombinant haplotypes distributed among 643 RILs in wheat-UK8, which is equivalent to 27.66 distinct recombinant haplotypes per RIL. Similarly, there are 17,643 distinct recombinant haplotypes distributed among 910 RILs in wheat-DE8, which is equivalent to 19.39 distinct recombinant haplotypes per RIL. When the interval is set to 10 cM, the counts and proportions of unique recombinant haplotypes decrease and the differences between wheat-UK8 and wheat-DE8 holds (Figure S3).

**Figure 3.**
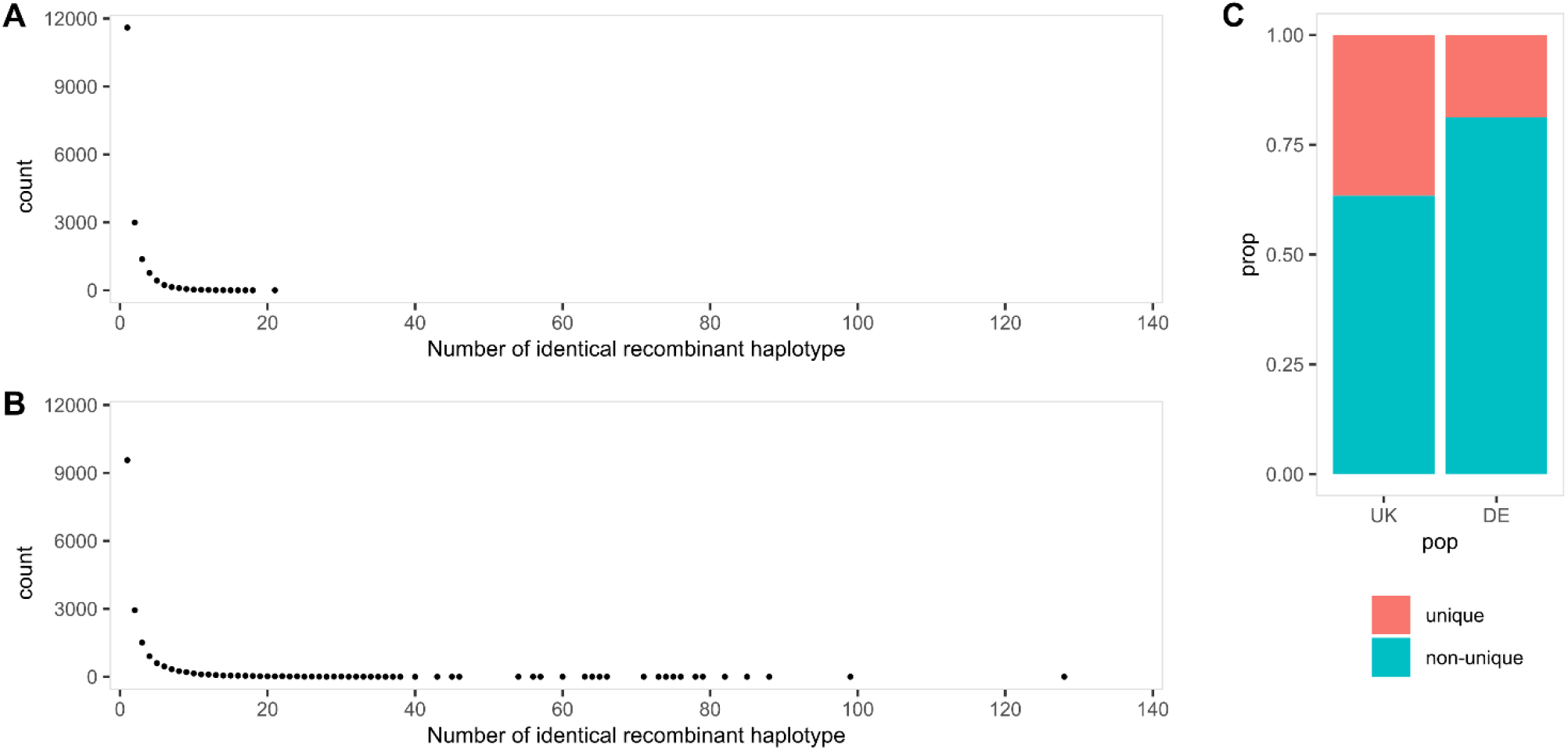
Distributions of unique and identical recombinant haplotypes in two wheat MAGIC populations. Recombinant haplotypes are considered identical if they are of the same founder pairs and present in the same 1 cM interval, otherwise unique. [**A**] Counts of the number of identical recombinant haplotypes in wheat-UK8. The left most point is the count of unique recombinant haplotypes. [**B**] Counts of the number of identical recombinant haplotypes in wheat-DE8. [**C**] Proportions of unique and non-unique (identical) recombinant haplotypes in wheat-UK8 and wheat-DE8.

**Table 3.** Five MAGIC population designs tested in magicdesign. The numbers of replicates, selfing generations and crosses are listed separately for each generation. For example, in design 1, the eight-way individuals are replicated three times and then selfed for four generations. The total number of crosses is shown in parentheses.

|  | Design 1 | Design 2 | Design 3 | Design 4 | Design 5 |
| --- | --- | --- | --- | --- | --- |
| Founders | 8 | 8 | 8 | 8 | 8 |
| Type | Full | Partial<br>balanced | Partial<br>unbalanced | Partial<br>balanced | Basic |
| Replicates | 1, 1, 3 | 1, 9, 15 | 1, 9, 15 | 1, 9, 15 | 1, 4, 4, 15 |
| Selfing | 0, 0, 4 | 0, 0, 4 | 0, 0, 4 | 0, 1, 4 | 0, 0, 0, 4 |
| Crosses | 28, 210, 315<br>(553) | 28, 14, 63<br>(105) | 19, 13, 63<br>(95) | 28, 14, 63<br>(105) | 4, 2, 4, 64<br>(74) |
| Generations | 7 | 7 | 7 | 8 | 8 |
| RIL | 945 | 945 | 945 | 945 | 960 |
| Funnel | 315 | 7 | 7 | 7 | 1 |

### Empirical evaluation of three other MAGIC populations

While not directly comparable, the relationship between MAGIC population designs and the distributions of recombinant haplotypes in three other datasets remains consistent. Similar to wheat-DE8, the cowpea and tomato MAGIC populations were created from a basic design and thus have a skewed distribution of recombinant haplotypes (Figure 4). The recombinant haplotypes from two-way founder pairs are higher than the other recombinant haplotypes. In cowpea, the two-way recombinant haplotypes are 0.936 ± 0.171 (mean ± standard deviation) per RIL while the other recombinant haplotypes are 0.384 ± 0.123 per RIL. In tomato, the two-way recombinant haplotypes are 0.907 ± 0.095 per RIL and the other recombinant haplotypes are 0.416 ± 0.116 per RIL. On the other hand, wheat-UK16 was created from a partial balanced design and does not have any skew in its distribution of recombinant haplotypes (Figure S4). The recombinant haplotypes are 0.878 ± 0.102 per RIL.

**Figure 4.**
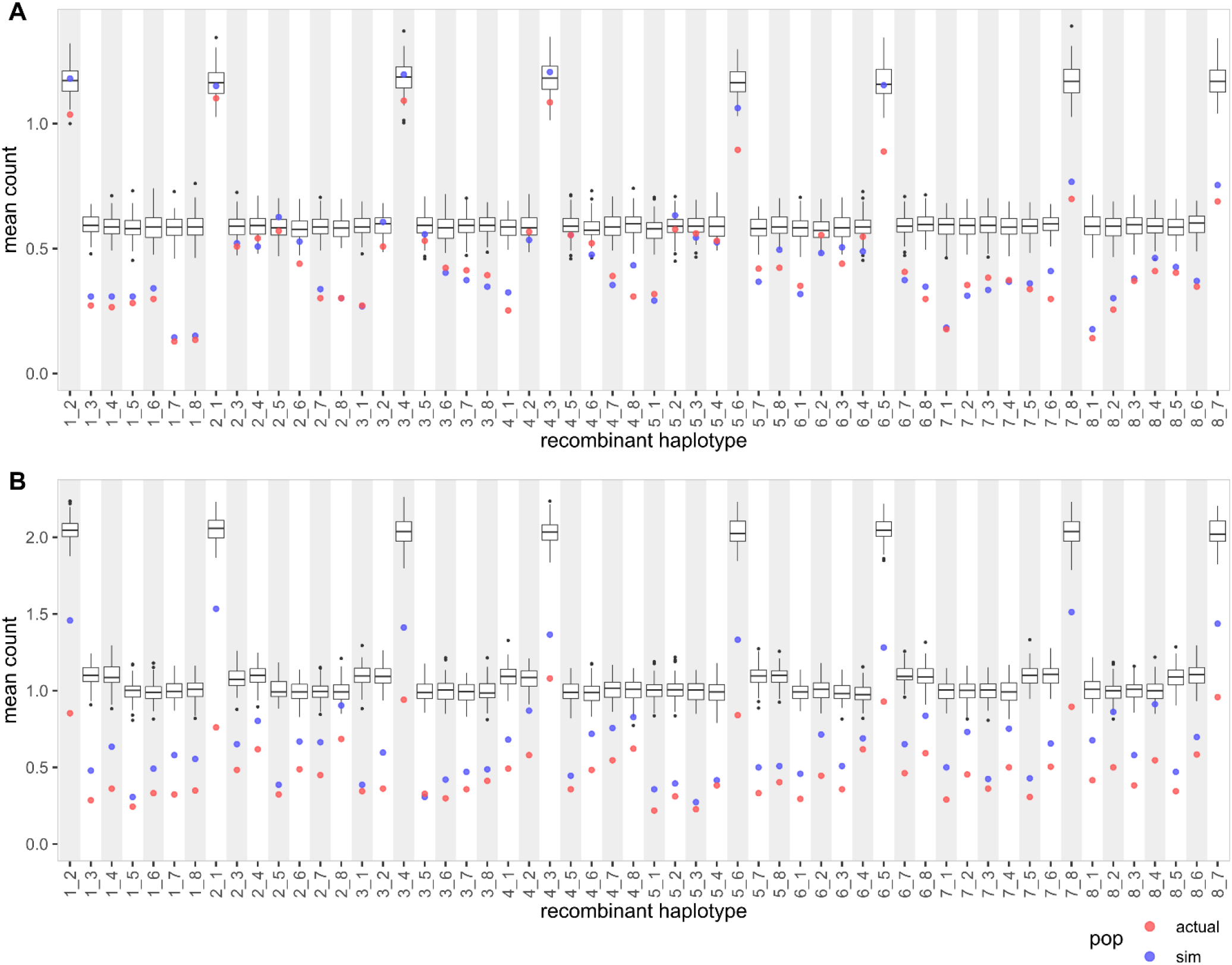
Distributions of recombinant haplotypes in cowpea and tomato MAGIC populations. [**A**] Plot shows mean count of each recombinant haplotype in a single RIL in cowpea. The boxplot shows mean count from true founder genotypes (100 simulated iterations). The red and blue points show mean count from inferred founder genotypes. [**B**] Plot shows mean count of each recombinant haplotype in a single RIL in tomato.

### Minimum probability for calling founder genotype

The minimum probability for calling founder genotype is important for the identification of recombinant haplotypes, and our simulation results suggest that the range of 0.4 to 0.6 gives a good balance of correct, incorrect and missing founder genotype calls (Figure 5A and 5B). The results are similar between simulated wheat-UK8 and wheat-DE8 populations, so only results from the simulated wheat-UK8 population are elaborated here. At a minimum probability of 0.4, the correct, incorrect and missing founder calls are 69%, 16% and 15% of the total markers, respectively. At a minimum probability of 0.5, the rates are 64%, 11% and 25%. At a minimum probability of 0.6, the rates are 58%, 6% and 36%. As the minimum probability increases, the rates of correct and incorrect founder calls decrease while the missing rate increases. In order to avoid the issue of having two or more founder probabilities above the threshold, the minimum probability can be set to 0.5 or higher.

**Figure 5.**
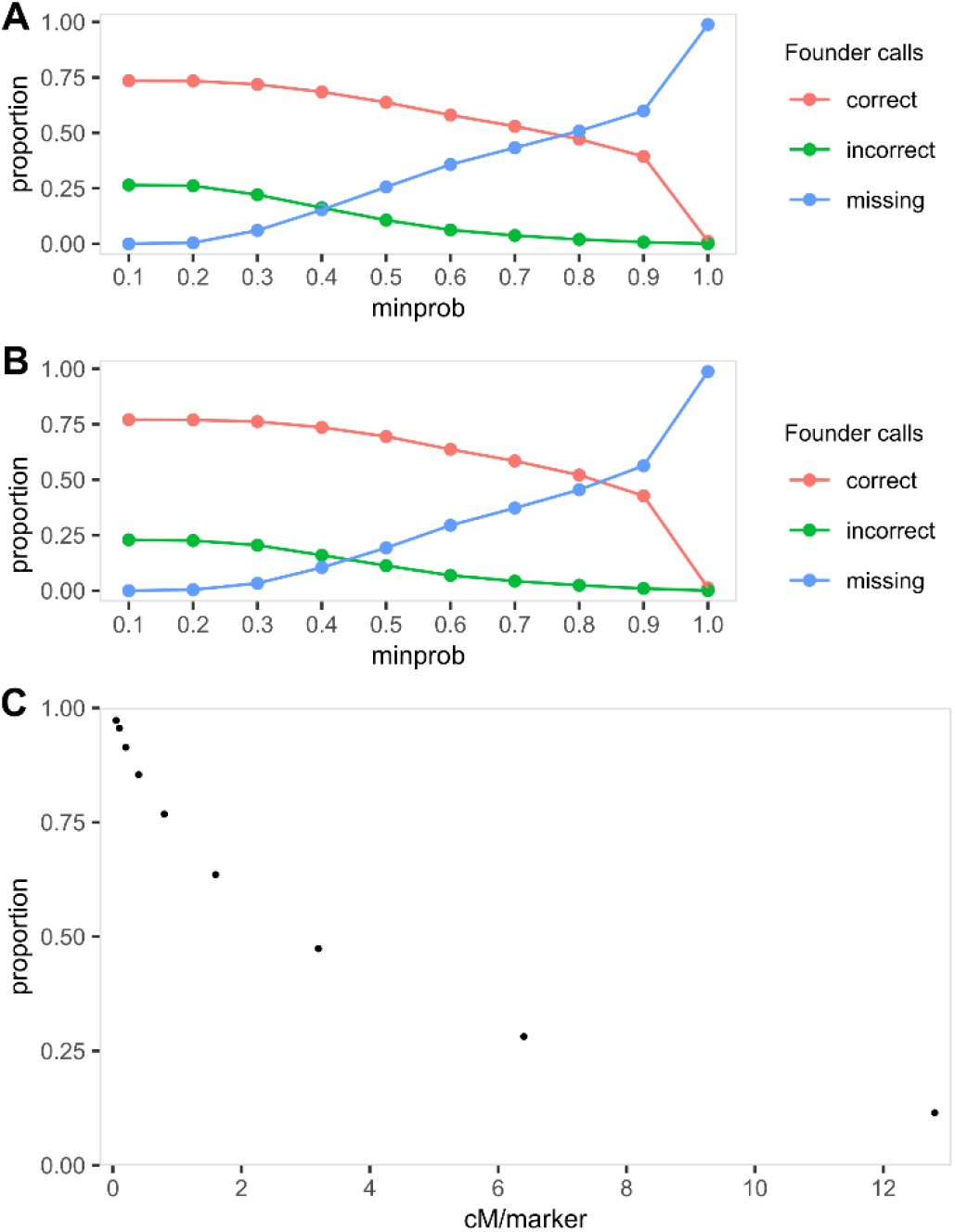
Ideal threshold for inferring founder genotypes and marker density in MAGIC population. **[A]** Proportions of correct, incorrect and missing founder genotypes inferred at different minimum probability (minprob) in simulated wheat-UK8 population. [**B**] Proportions of correct, incorrect and missing founder genotypes inferred at different minprob in simulated wheat-DE8 population. [**C**] Proportions of recombinant haplotypes recovered (PRHR) at different marker density along a simulated chromosome of 200 cM. The marker density is adjusted by having markers equally spaced at 0.05, 0.10, 0.20, 0.40, 0.80, 1.60, 3.20, 6.40 and 12.80 cM apart.

### Marker density in MAGIC population

In all five analyzed datasets, the proportion of recombinant haplotypes recovered (PRHR) is higher in populations genotyped at higher marker density (Table 1). PRHR is computed by taking the number of recombinant haplotypes in actual dataset divided by the true number of recombinant haplotypes in simulated dataset. In wheat-UK8 and wheat-DE8 with an average marker distance of 1.024 cM, the PRHR is approximately one-third (Figure 2). Even though the tomato population is genotyped at a lower marker density with an average marker distance of 1.603 cM, the PRHR is higher than in the two wheat populations (Figure 4B). This is likely because the markers on the wheat D-genome are generally sparse due to its low diversity (Akhunov et al. 2010). The cowpea population is genotyped at a high marker density with an average marker distance of 0.030 cM, in which the PRHR is approximately two-thirds (Figure 4A). Lastly, wheat-UK16 is genotyped at the highest marker density of all analyzed datasets with an average marker distance of 0.005 cM, and it has the highest PRHR of almost 80% (Figure S4). Marker density is an important factor in identifying fine-scale recombination breakpoints in MAGIC populations.

Under ideal conditions, where the markers are evenly spaced, a marker distance of 0.20 cM or less between two adjacent markers is sufficient to achieve a PRHR of at least 90% (Figure 5C). We tested recombinant haplotype recovery rates for markers that are evenly spaced across 0.05, 0.10, 0.20, 0.40, 0.80, 1.60, 3.20, 6.40 and 12.80 cM. At the smallest tested distance of 0.05 cM, approximately 97% of the true recombinant haplotypes can be recovered. As the distance increases, the recovery rate decreases. At the largest tested distance of 12.80 cM, approximately 11% of the true recombinant haplotypes can be recovered. These results are more optimistic than the actual results (Table 1). In practice, more markers are required to achieve the same recovery rate for any given marker density because markers are not evenly distributed across the whole genome. In addition, the discrepancy between simulated and actual results can also be attributed to marker quality. For example, the markers on the wheat D-genomes are generally sparser than on the others.

### magicdesign: a tool to create and test MAGIC population designs

Given that MAGIC population construction requires a lot of time and effort, and that design choices can impact population attributes, there is a need for a “free trial” before committing to create a MAGIC population. Here, we introduce an R package called “magicdesign”, which is specifically made for creating and testing various MAGIC population designs via simulation. Alternatively, we also provide a user-friendly Shiny app version called “magicdesignee” which implements the “magicdesign” R package in its back-end. Therefore, minimal R knowledge required for users to use “magicdesignee”.

Briefly, the “magicdesign” package workflow can be described as: (1) design creation, (2) population simulation, and (3) comparative analysis. In the design creation step, the package creates a crossing scheme that spans from the founders to the final RILs based on user inputs. In the population simulation step, the package simulates a MAGIC RIL population constructed from the crossing scheme, and repeats over multiple iterations. At this point, the first two steps may be repeated for other MAGIC population designs. Finally, in the comparative analysis step, the package extracts information from previously tested designs and summarizes the results illustratively. Additional details on each step are described in subsequent sections.

### Design creation

In a structured design, the design creation step takes various user inputs to create a crossing scheme. The major inputs include number of founders, number of funnels or funnel sets, and a balanced design indicator. Based on how these inputs are specified, one of the structured designs (Full, Partial Balanced, Partial Unbalanced, Basic) as shown in Figure 1A is created. As defined previously, a balanced design has an equal number of founders among the funnels and equal frequency of founder pairs at each crossing generation. This design creation step works for either power of 2 (P2) or non-power of 2 (NP2) number of founders. Currently, the allowed range of number of founders is any integer between 3 and 128. The allowed number of funnels or funnel sets varies according to the number of founders and the balanced design indicator, and the full list is provided in Table S1.

Finding a balanced design requires more computation power than finding an unbalanced design. This is because the balanced design requires many funnel permutations to be evaluated while the unbalanced design randomly sample the required number of funnel permutations. To reduce the computational burden, we have identified alternative methods that are less computationally intensive. In the case of 8 founders, we have searched through all 315^7^ possible combinations and identified 720 partial balanced funnel sets. Each of these partial balanced funnel set has 7 funnels, and can be combined with another non-overlapping partial balanced funnel set to form a larger partial balanced funnel set. In the case of 16 founders, the number of possible combinations is very large and so we opted for a different approach. To start, we obtained the 15 funnels from Scott et al. (2021), which is a partial balanced set for 16 founders. We searched through all 3^15^ possible permutations of eight- and sixteen-way crosses in these funnels and identified 7,776 partial balanced funnel sets. More partial balanced funnel sets could be found by searching through all 315^15^ possible permutations of four-way crosses, however, that was beyond our available computational capacity. Unlike the case of 8 founders, these funnel sets do overlap and thus cannot be combined to form a larger set. Instead, by randomly swapping the founders from a starting partial balanced funnel set, a non-overlapping partial balanced funnel set can be created and merged to form a larger set. For other numbers of founders between 4 and 16, the balanced design is created based on a nested incomplete block design (NIBD) generated using the “blocksdesign” package (Edmondson 2020; Edmondson 2021). A MAGIC funnel is analogous to a NIBD as the founders in two-way crosses (experimental block of two plots) are nested within four-way crosses, founders in four-way crosses are nested within eight-way crosses, and so on. Currently, a balanced design in “magicdesign” is limited to 16 or less founders as there is not yet an efficient method for larger number of founders.

In addition, “magicdesign” provides options to further modify the MAGIC population design by specifying the number of replicates, number of selfing generations, and an additional crossing indicator. The number of replicates determines how many seeds from a cross are retained. This can help to increase the haplotype diversity in the MAGIC population when the seeds are not genetically identical. In the case of inbred founders, replicates of two-ways individuals are all identical but not replicates of four-ways (or higher) individuals. The number of selfing generations determine how many generations of selfing are required after each cross. Typically, the selfing step is only applied after the last crossing generation as a way to reduce heterozygosity in the RILs. However, selfing prior to that may be beneficial in increasing recombinant haplotypes. Lastly, the additional crossing indicator allows for an extra crossing generation to further increase recombinant haplotypes. This is similar to the approach taken by Stadlmeier et al. (2018) and Shah et al. (2019).

Alternatively, any MAGIC population design that is not available directly in “magicdesign” can be created by supplying a complete pedigree. The only requirement for the pedigree is that it must detail all crosses involved from the founders to the final RILs. This option provides a greater flexibility to accommodate for semi-structured or unstructured designs. Furthermore, it is also possible to modify a design created from “magicdesign” and provide the pedigree of the modified design.

### Population simulation

Once a MAGIC population design is created, “magicdesign” simulates a population based on the design and other user inputs. The major inputs include distance between markers, chromosome genetic lengths, number of simulations and recombinant haplotype interval size. The simulation step will create evenly-spaced markers based on the distance between markers and chromosome genetic lengths. All founders are considered unique and so each of these markers is used to encode for the founder genotypes. The desired number of simulations is selected. In addition, the recombinant haplotype interval size determines the distance between two markers to look for recombinant haplotypes.

### Comparative analysis

After simulating one or more designs, the final step is to compare the design qualities in terms of recombinant haplotype proportions and distribution of founder genomes in the RILs. In general, a good MAGIC population design should yield consistently higher recombinant haplotype proportions as well as an even distribution of founder genomes compared to other designs.

To demonstrate comparative analysis with “magicdesign”, the five designs in Table 3 are used as examples. These designs are all applied to a fictitious species with five chromosomes of 1.0, 1.5, 2.0, 2.5, 3.0 centiMorgans (cM) length. All five designs are created based on a MAGIC population of 8 founders. Design 1 is a full design and so it has all 315 funnels. Design 2 and 4 are both partial balanced design with one funnel set (7 funnels), and the only difference between them is that the four-way individuals in design 4 are selfed once before making eight-way crosses. Design 3 is similar to design 2 except it is a partial unbalanced design with 7 funnels. Lastly, design 5 is a basic design with 1 funnel inspired by the design used in Stadlmeier et al. (2018). The numbers of replicates are varied for each design to achieve similar final RIL population size close to 1,000. Aside from design 1 which has the highest number of crosses, the other designs have fairly similar numbers of crosses. Design 1, 2 and 3 require 7 generations from founders to RILs, while design 4 and 5 require 8 generations because of the additional selfing and crossing generation respectively.

First, we investigated the designs’ effects on recombinant haplotypes within a 5 cM interval. In term of total recombinant haplotypes, a good design should have high mean with low variance. For design 1 to 5 respectively, the means are 0.167, 0.167, 0.169, 0.186 and 0.202 while the variances are 0.000158, 0.000244, 0.000293, 0.000452 and 0.003000 (Figure 6A). The means are similar in design 1, 2 and 3, slightly higher in design 4 and highest in design 5. However, the variances are lowest in design 1, similar in design 2 and 3, slightly larger in design 4, and substantially larger in design 5.

**Figure 6.**
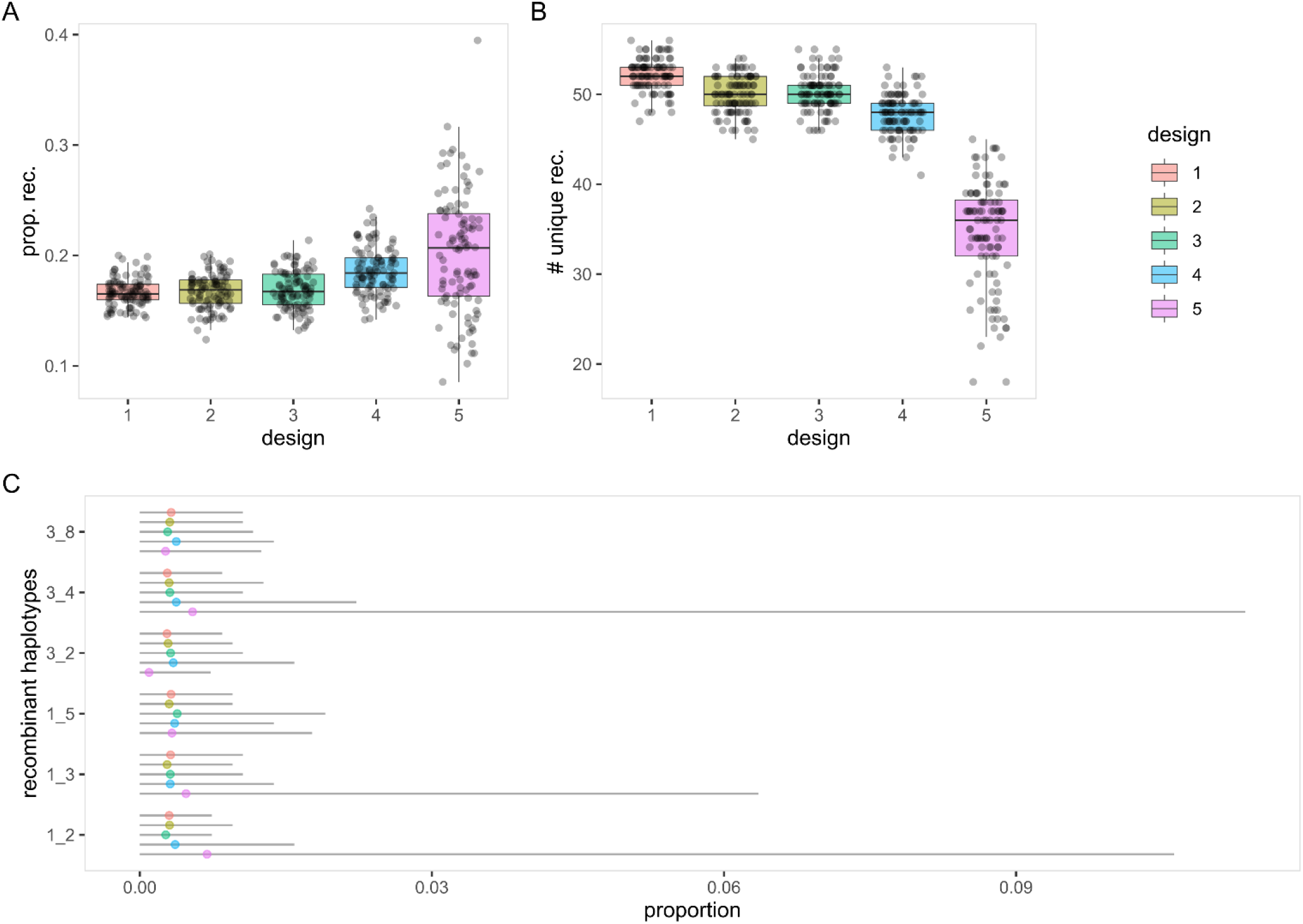
Distributions of recombinant haplotypes in five MAGIC population designs. Recombinant haplotypes are evaluated within a 5 cM interval over 100 iterations of simulation. [**A**] Proportions of total recombinant haplotypes. [**B**] Number of unique recombinant haplotypes. [**C**] Proportions of six chosen recombinant haplotypes. Complete results are available in Table S5.

In any RIL derived from an 8-founder MAGIC population, there are 56 distinct recombinant haplotypes and a good design should have high mean with low variance. The means for the number of unique recombinant haplotypes are 52.20, 49.93, 50.29, 47.81 and 34.51 for design 1 to 5 respectively while the variances are 3.31, 4.29, 4.21, 4.68 and 34.78 for design 1 to 5 respectively (Figure 6B). The means are highest in design 1, similar in design 2 and 3, slightly lower in design 4 and lowest in design 5. The variances follow a similar but reverse trend as the means except for design 5 where the variance is over seven times greater. Equivalently, the coefficients of variation (CVs) are 0.035, 0.041, 0.041, 0.045 and 0.171 for design 1 to 5 respectively. That for design 5 is approximately four times that for the other designs.

The distributions of individual recombinant haplotype should be consistent across all recombinant haplotypes with minimal variability across simulations in a good design. With the exception of design 5, all other designs have similar distributions of individual recombinant haplotype (Figure 6C and Table S5). Similar to wheat-DE8 (Figure 2B), cowpea and tomato (Figure 4), design 5 has more two-ways recombinant haplotypes than other recombinant haplotypes. Furthermore, the spreads of two-ways recombinant haplotypes in design 5 are much higher than the others, which imply low consistency.

Similar to the previous criterion, the proportions of founder genomes should be consistent across all founders with low variability across simulations in a good design. With 8 founders, the expected proportion of each founder genome in a population is 0.125. The proportions are within 0.01 of expectation for all designs except for design 5, which has 5 out of 8 proportions exceeding the range (Figure 7A). On the other hand, the variances are lowest in design 1, slightly higher in design 2, 3 and 4, and highest in design 5 (Figure 7A).

**Figure 7.**
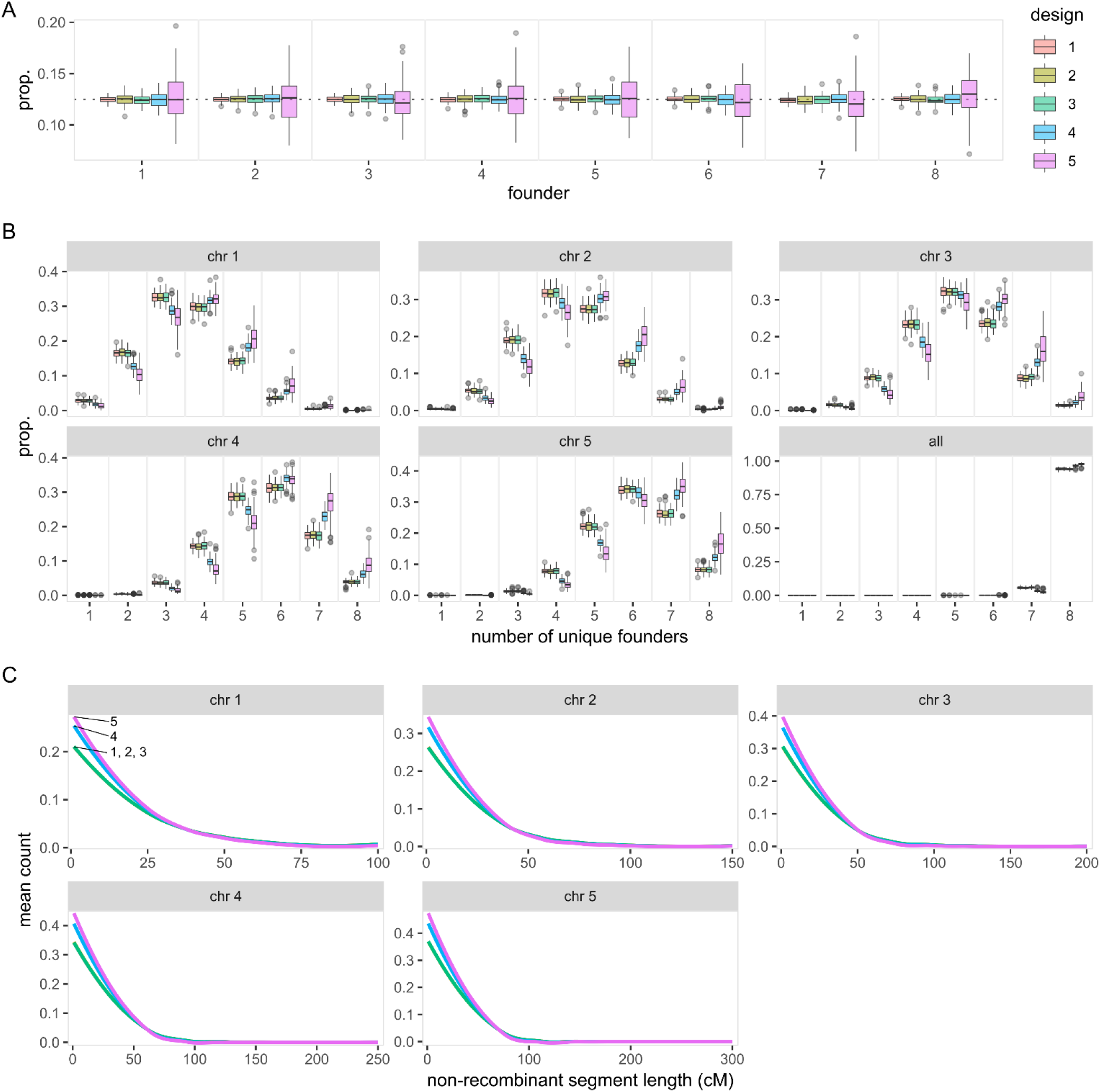
Distributions of founder genomes in five MAGIC population designs. Founder genomes are evaluated from 100 iterations of simulation. [**A**] Proportions of each founder genome in the MAGIC RILs. [**B**] Proportions of the MAGIC RILs carrying tracts of 1 to 8 unique founder genomes in each chromosome. [**C**] Mean count of non-recombinant segment length in each RIL’s chromosome.

In any single chromosome, a RIL can carry tracts of 1 to 8 unique founder genomes and it is generally better to have more unique founder genomes. In the shortest chromosome (chromosome 1), design 1, 2 and 3 frequently produce 3 unique founders, while design 4 and 5 frequently produce 4 unique founders (Figure 7B). In the longest chromosome (chromosome 5), design 1, 2, 3 and 4 frequently produce 6 unique founders while design 5 frequently produces 7 unique founders (Figure 7B).

Lastly, a good design should have short non-recombinant segments. Across all chromosomes, design 5 has the most short non-recombinant segments, followed by design 4, and design 1, 2 and 3 being undiscernible (Figure 7C).

Of all the five designs considered here, each has its own advantages and disadvantages. Design 1, 2 and 3 are highly similar except that design 1 tends to show smaller variability than the other two at the cost of more crossing work required. Design 4 is slightly better than the first three in most occasions, although it is slightly more variable and requires one additional generation. Design 5 is generally poor and should be avoided if possible, although the additional crossing generation helps in increasing the number of unique founders and reducing non-recombinant segment lengths. Of all designs considered, design 4 appears to be the best option if the additional generation is acceptable, otherwise either design 2 or 3 is a good option too. Across all the metrics used for comparisons in “magicdesign”, there is no observable difference between design 2 (balanced) and 3 (unbalanced).

## Discussion

Ease of design appears as a major factor in driving the design choices in currently available MAGIC populations. These MAGIC populations are predominantly made of 4 or 8 founders crossed using a basic design (Figure 1B). There are several possible explanations to the choice popularity. First, P2 designs are easier to handle than NP2 designs since the individuals in every funnel are halved at every crossing generation. Besides, 4 and 8 founders are effectively the lowest numbers of founders available in P2 designs, and higher numbers of founders require more generations of crossing and may increase the design complexity. Of all the explored designs, the basic design likely requires the least amount of effort in population construction. The only other option that may rival a basic design is the unstructured design with random mating, which often relies on segregation of male sterility loci. Unfortunately, this system is not always readily available in every species and may restrict the founder choices.

Choice of MAGIC population design plays a critical role in determining the recombination landscape in the RILs. In the comparison between wheat-UK8 and wheat-DE8, we identified a bias in individual recombinant haplotypes in the basic design but not the partial design (Figure 2). The bias resulted in more two-way recombinant haplotypes than other recombinant haplotypes. The bias might be exacerbated if the pairs of founders in two-ways are genetically more similar than others, which can happen if the founders stratify into two or more groups. It is possible to avoid pairing the founders of the same groups in two-ways if the grouping is known. For example, Pascual et al. (2015) made the two-way crosses by crossing tomato founders with large fruits to founders with small fruits, and Ogawa et al. (2018) followed similarly by crossing *indica* rice founders to *japonica* rice founders. This countermeasure is only possible if the numbers of founders are equal between groups, but not if the founders cannot be subdivided equally like the barley (Sannemann et al. 2015), cowpea (Huynh et al. 2018) and wheat (Stadlmeier et al. 2018) MAGIC populations.

In addition to the bias, the basic design also resulted in a lower proportion of unique recombinant haplotypes than the partial design (Figure 3). Since a basic design always has less funnels than any other designs, high replication of cross progeny is required to bring the number of RILs up. In general, replicates reduce the amount of crossing work required in prior generation by keeping more than one progeny from a single cross to advance. The recombination landscape in these replicated individuals is non-independent because any prior recombinations are passed down from their parents. The detriments from replication can be minimized by replicating in earlier generations as subsequent crosses will reduce the non-independence among replicates. In a MAGIC population with 8 inbred founders, the earliest meaningful replication would be the four-way individuals. However, replicates prior to the final crosses do increase the amount of downstream crossing work, and so it is important to consider the trade-offs between available work resources and uniqueness of recombinant haplotypes.

High marker density is needed to capture the highly recombined genomes of MAGIC RILs. We used the proportion of recombinant haplotypes recovered (PRHR) as a measure of how well the markers capture recombinant haplotypes. PRHRs in the five analyzed datasets correlate well with the marker density. Even with the high marker density in wheat-UK16, the PRHR is only 0.799 (Table 1), which suggests that one-fifth of the recombinant haplotypes is still missing. Some explanations include the sparser marker density in the D-genomes, uneven marker density along the genomes and segregation distortions of the founder genomes. To generalize the relationship between marker density and recombinant haplotypes further, our simulation results showed that marker distance of 0.80 cM or less is sufficient to recover over three-quarters of the recombinant haplotypes. Despite the results from actual datasets being less optimistic than the results from simulation, the importance of high marker density in MAGIC populations still holds.

Given that the advantages and disadvantages of different MAGIC population designs are largely unexplored, the “magicdesign” package serves as an important tool to create and test different designs. Specifically, “magicdesign” provides the opportunity to evaluate the options before committing to years of effort in constructing MAGIC populations. In our examples, an additional selfing generation offers a simple path to improvement (Figure 6 and 7), especially in inbreeding species. When used in combination with speed breeding (Watson et al. 2018), the additional time due to selfing can be minimized. In addition, “magicdesign” also acts as a bridging tool for researchers who are new to MAGIC populations by providing a starting point to creating a MAGIC population. The opportunity to create and test different designs will encourage innovation in MAGIC population designs rather than relying on previously used designs. Overall, “magicdesign” is a valuable resource for unifying the process of creating and testing MAGIC population designs, and providing the flexibility for additional features to be included in future updates as the package grows.

## Acknowldgements

We thank Rajiv Sharma, Ian Dawson and David Marshall for helpful discussion. We also thank the UK Crop Diversity group (https://www.cropdiversity.ac.uk/) for providing the High Performance Computing (HPC) resource used in evaluating large permutations.

## Supplementary Materials

### Supplementary Text

#### Source datasets

Links to source datasets without DOI have been archived at https://web.archive.org on April 13, 2021.

- wheat-UK8: https://www.niab.com/research/agricultural-crop-research/resources/niab-magic-population-resources
- wheat-DE8: https://doi.org/10.1186/s12864-018-4915-3 (Table S1)
- cowpea: https://doi.org/10.1111/tpj.13827 (Data S1)
- tomato: https://doi.org/10.1111/pbi.12282 (Table S1 and S2)
- wheat-UK16: http://mtweb.cs.ucl.ac.uk/mus/www/MAGICdiverse/

## Supplementary Tables

**Table S1.** Number of funnels in a partial balanced set or full design. In P2 design, a balanced design is defined where all founders are present equally in the funnels and all founder pairings are present equally at each level of crosses. The number of funnels required for a partial balanced set is *n* − 1 and for full design is *n*!/2^*n*−1^. In NP2 design, a balanced design is less strictly defined where only all founders are present equally in the funnels, without the latter requirement in P2. The number of funnels required for a partial balanced set is *x* where *x* must be the smallest positive integer that satisfies (*x* · *n*_0_)⁄*n mod* 1 = 0 and *n*_0_ is the next highest P2 *n*. Number of funnels in a full design is provided up to *n* = 8 as the number of funnels in a full design for higher *n* is impractical.

| n | set | full | n | set | full | n | set | full | n | set | full |
| --- | --- | --- | --- | --- | --- | --- | --- | --- | --- | --- | --- |
| 3 | 3 | 3 | 35 | 35 | NA | 67 | 67 | NA | 99 | 99 | NA |
| 4 | 3 | 3 | 36 | 9 | NA | 68 | 17 | NA | 100 | 25 | NA |
| 5 | 5 | 240 | 37 | 37 | NA | 69 | 69 | NA | 101 | 101 | NA |
| 6 | 3 | 855 | 38 | 19 | NA | 70 | 35 | NA | 102 | 51 | NA |
| 7 | 7 | 945 | 39 | 39 | NA | 71 | 71 | NA | 103 | 103 | NA |
| 8 | 7 | 315 | 40 | 5 | NA | 72 | 9 | NA | 104 | 13 | NA |
| 9 | 9 | NA | 41 | 41 | NA | 73 | 73 | NA | 105 | 105 | NA |
| 10 | 5 | NA | 42 | 21 | NA | 74 | 37 | NA | 106 | 53 | NA |
| 11 | 11 | NA | 43 | 43 | NA | 75 | 75 | NA | 107 | 107 | NA |
| 12 | 3 | NA | 44 | 11 | NA | 76 | 19 | NA | 108 | 27 | NA |
| 13 | 13 | NA | 45 | 45 | NA | 77 | 77 | NA | 109 | 109 | NA |
| 14 | 7 | NA | 46 | 23 | NA | 78 | 39 | NA | 110 | 55 | NA |
| 15 | 15 | NA | 47 | 47 | NA | 79 | 79 | NA | 111 | 111 | NA |
| 16 | 15 | NA | 48 | 3 | NA | 80 | 5 | NA | 112 | 7 | NA |
| 17 | 17 | NA | 49 | 49 | NA | 81 | 81 | NA | 113 | 113 | NA |
| 18 | 9 | NA | 50 | 25 | NA | 82 | 41 | NA | 114 | 57 | NA |
| 19 | 19 | NA | 51 | 51 | NA | 83 | 83 | NA | 115 | 115 | NA |
| 20 | 5 | NA | 52 | 13 | NA | 84 | 21 | NA | 116 | 29 | NA |
| 21 | 21 | NA | 53 | 53 | NA | 85 | 85 | NA | 117 | 117 | NA |
| 22 | 11 | NA | 54 | 27 | NA | 86 | 43 | NA | 118 | 59 | NA |
| 23 | 23 | NA | 55 | 55 | NA | 87 | 87 | NA | 119 | 119 | NA |
| 24 | 3 | NA | 56 | 7 | NA | 88 | 11 | NA | 120 | 15 | NA |
| 25 | 25 | NA | 57 | 57 | NA | 89 | 89 | NA | 121 | 121 | NA |
| 26 | 13 | NA | 58 | 29 | NA | 90 | 45 | NA | 122 | 61 | NA |
| 27 | 27 | NA | 59 | 59 | NA | 91 | 91 | NA | 123 | 123 | NA |
| 28 | 7 | NA | 60 | 15 | NA | 92 | 23 | NA | 124 | 31 | NA |
| 29 | 29 | NA | 61 | 61 | NA | 93 | 93 | NA | 125 | 125 | NA |
| 30 | 15 | NA | 62 | 31 | NA | 94 | 47 | NA | 126 | 63 | NA |
| 31 | 31 | NA | 63 | 63 | NA | 95 | 95 | NA | 127 | 127 | NA |
| 32 | 31 | NA | 64 | 63 | NA | 96 | 3 | NA | 128 | 127 | NA |
| 33 | 33 | NA | 65 | 65 | NA | 97 | 97 | NA |  |  |  |
| 34 | 17 | NA | 66 | 33 | NA | 98 | 49 | NA |  |  |  |

**Table S2.**
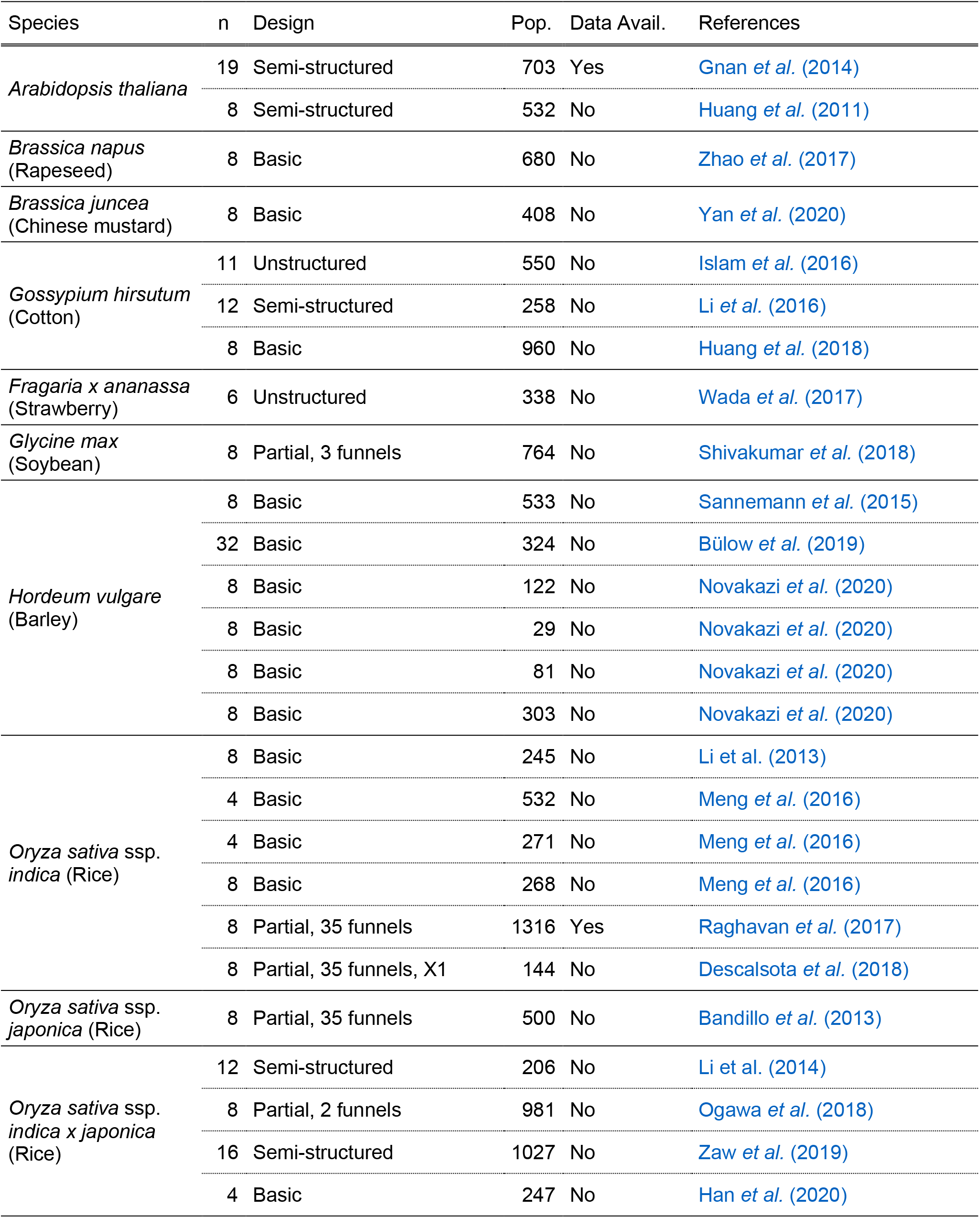
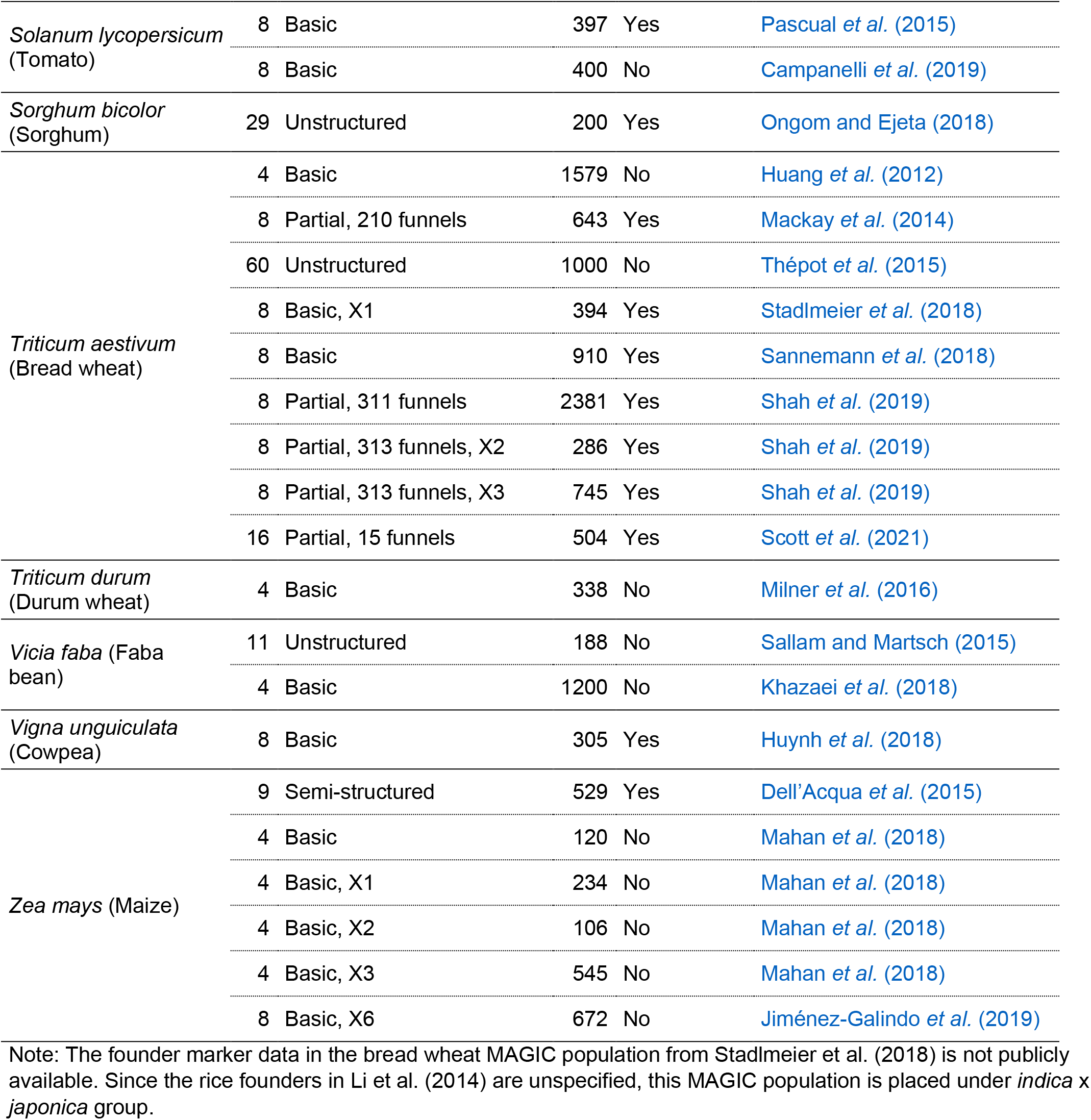
Published MAGIC populations. Most, if not all, of the MAGIC populations that have been described in either published literature or pre-prints are listed as of March 17, 2021.

| Species | n | Design | Pop. | Data Avail. | References |
| --- | --- | --- | --- | --- | --- |
| *Arabidopsis thaliana* | 19 | Semi-structured | 703 | Yes | Gnan et al. (2014) |
|  | 8 | Semi-structured | 532 | No | Huang et al. (2011) |
| *Brassica napus*<br>(Rapeseed) | 8 | Basic | 680 | No | Zhao et al. (2017) |
| *Brassica juncea*<br>(Chinese mustard) | 8 | Basic | 408 | No | Yan et al. (2020) |
| *Gossypium hirsutum*<br>(Cotton) | 11 | Unstructured | 550 | No | Islam et al. (2016) |
|  | 12 | Semi-structured | 258 | No | Li et al. (2016) |
|  | 8 | Basic | 960 | No | Huang et al. (2018) |
| *Fragaria x ananassa*<br>(Strawberry) | 6 | Unstructured | 338 | No | Wada et al. (2017) |
| *Glycine max*<br>(Soybean) | 8 | Partial, 3 funnels | 764 | No | Shivakumar et al. (2018) |
| *Hordeum vulgare*<br>(Barley) | 8 | Basic | 533 | No | Sannemann et al. (2015) |
|  | 32 | Basic | 324 | No | Bülow et al. (2019) |
|  | 8 | Basic | 122 | No | Novakazi et al. (2020) |
|  | 8 | Basic | 29 | No | Novakazi et al. (2020) |
|  | 8 | Basic | 81 | No | Novakazi et al. (2020) |
|  | 8 | Basic | 303 | No | Novakazi et al. (2020) |
| *Oryza sativa* ssp.<br>*indica* (Rice) | 8 | Basic | 245 | No | Li et al. (2013) |
|  | 4 | Basic | 532 | No | Meng et al. (2016) |
|  | 4 | Basic | 271 | No | Meng et al. (2016) |
|  | 8 | Basic | 268 | No | Meng et al. (2016) |
|  | 8 | Partial, 35 funnels | 1316 | Yes | Raghavan et al. (2017) |
|  | 8 | Partial, 35 funnels, X1 | 144 | No | Descalsota et al. (2018) |
| *Oryza sativa* ssp.<br>*japonica* (Rice) | 8 | Partial, 35 funnels | 500 | No | Bandillo et al. (2013) |
| *Oryza sativa* ssp.<br>*indica x japonica*<br>(Rice) | 12 | Semi-structured | 206 | No | Li et al. (2014) |
|  | 8 | Partial, 2 funnels | 981 | No | Ogawa et al. (2018) |
|  | 16 | Semi-structured | 1027 | No | Zaw et al. (2019) |
|  | 4 | Basic | 247 | No | Han et al. (2020) |
| *Solanum lycopersicum*<br>(Tomato) | 8 | Basic | 397 | Yes | Pascual et al. (2015) |
|  | 8 | Basic | 400 | No | Campanelli et al. (2019) |
| *Sorghum bicolor*<br>(Sorghum) | 29 | Unstructured | 200 | Yes | Ongom and Ejeta (2018) |
| *Triticum aestivum*<br>(Bread wheat) | 4 | Basic | 1579 | No | Huang et al. (2012) |
|  | 8 | Partial, 210 funnels | 643 | Yes | Mackay et al. (2014) |
|  | 60 | Unstructured | 1000 | No | Thépot et al. (2015) |
|  | 8 | Basic, X1 | 394 | Yes | Stadlmeier et al. (2018) |
|  | 8 | Basic | 910 | Yes | Sannemann et al. (2018) |
|  | 8 | Partial, 311 funnels | 2381 | Yes | Shah et al. (2019) |
|  | 8 | Partial, 313 funnels, X2 | 286 | Yes | Shah et al. (2019) |
|  | 8 | Partial, 313 funnels, X3 | 745 | Yes | Shah et al. (2019) |
|  | 16 | Partial, 15 funnels | 504 | Yes | Scott et al. (2021) |
| *Triticum durum*<br>(Durum wheat) | 4 | Basic | 338 | No | Milner et al. (2016) |
| *Vicia faba* (Faba bean) | 11 | Unstructured | 188 | No | Sallam and Martsch (2015) |
|  | 4 | Basic | 1200 | No | Khazaei et al. (2018) |
| *Vigna unguiculata*<br>(Cowpea) | 8 | Basic | 305 | Yes | Huynh et al. (2018) |
| *Zea mays* (Maize) | 9 | Semi-structured | 529 | Yes | Dell'Acqua et al. (2015) |
|  | 4 | Basic | 120 | No | Mahan et al. (2018) |
|  | 4 | Basic, X1 | 234 | No | Mahan et al. (2018) |
|  | 4 | Basic, X2 | 106 | No | Mahan et al. (2018) |
|  | 4 | Basic, X3 | 545 | No | Mahan et al. (2018) |
|  | 8 | Basic, X6 | 672 | No | Jiménez-Galindo et al. (2019) |

Note: The founder marker data in the bread wheat MAGIC population from Stadlmeier et al. (2018) is not publicly available. Since the rice founders in Li et al. (2014) are unspecified, this MAGIC population is placed under *indica* x *japonica* group.

**Table S3.** Number of informative recombinations in wheat-UK8 (full dataset). The number of informative recombinations (NR) is calculated for both simulated and actual wheat-UK8 dataset. Note: recombinant inbred line (RIL), Morgan (M).

| Chr | NR/RIL |  | NR/RIL/M |  |
| --- | --- | --- | --- | --- |
|  | sim | actual | sim | actual |
| 1A | 8.42 | 3.37 | 3.53 | 1.41 |
| 1B | 12.94 | 4.64 | 3.69 | 1.32 |
| 1D | 4.44 | 1.78 | 3.38 | 1.36 |
| 2A | 9.36 | 5.18 | 3.61 | 2.00 |
| 2B | 13.64 | 3.58 | 3.59 | 0.94 |
| 2D | 6.50 | 2.97 | 3.27 | 1.49 |
| 3A | 11.13 | 5.40 | 3.60 | 1.75 |
| 3B | 10.45 | 4.29 | 3.69 | 1.51 |
| 3D | 5.59 | 2.03 | 2.87 | 1.05 |
| 4A | 7.94 | 3.32 | 3.64 | 1.52 |
| 4B | 8.43 | 3.03 | 3.63 | 1.30 |
| 4D | 3.95 | 1.36 | 3.14 | 1.08 |
| 5A | 11.07 | 5.42 | 3.53 | 1.73 |
| 5B | 11.38 | 3.23 | 3.65 | 1.04 |
| 5D | 6.42 | 2.58 | 3.17 | 1.28 |
| 6A | 10.42 | 4.23 | 3.66 | 1.49 |
| 6B | 9.57 | 2.67 | 3.64 | 1.02 |
| 6D | 5.44 | 1.03 | 2.53 | 0.48 |
| 7A | 14.24 | 5.19 | 3.69 | 1.35 |
| 7B | 10.19 | 3.78 | 3.55 | 1.32 |
| 7D | 6.35 | 2.28 | 2.91 | 1.05 |
| All | 187.88 | 71.39 | 3.48 | 1.32 |

**Table S4.**
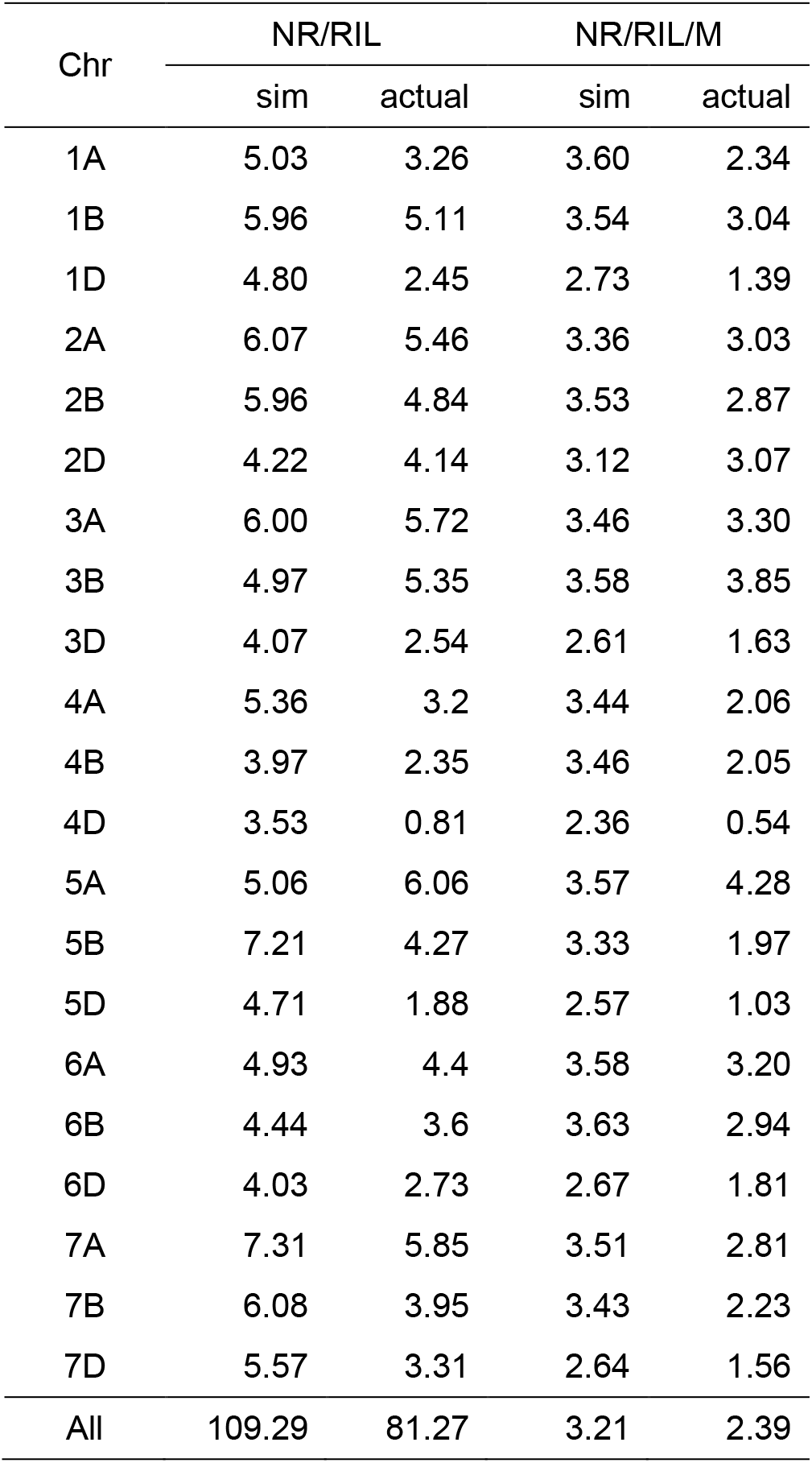
Number of informative recombinations in wheat-DE8 (full dataset). The number of informative recombinations (NR) is calculated for both simulated and actual wheat-DE8a dataset. Note: recombinant inbred line (RIL), Morgan (M).

**Table S5.** Proportions of individual recombinant haplotypes in five MAGIC population designs. The mean proportions (standard deviations in parentheses) are shown for each recombinant haplotype and design. Two-ways recombinant haplotypes in design 5 are highlighted.

| Rec. hap. | Design 1 | Design 2 | Design 3 | Design 4 | Design 5 |
| --- | --- | --- | --- | --- | --- |
| 1_2 | 0.0030 (0.0019) | 0.0031 (0.0020) | 0.0027 (0.0018) | 0.0036 (0.0036) | 0.0069 (0.0206) |
| 1_3 | 0.0032 (0.0021) | 0.0028 (0.0020) | 0.0031 (0.0022) | 0.0031 (0.0030) | 0.0048 (0.0108) |
| 1_4 | 0.0030 (0.0020) | 0.0029 (0.0025) | 0.0025 (0.0021) | 0.0033 (0.0028) | 0.0058 (0.0104) |
| 1_5 | 0.0032 (0.0019) | 0.0030 (0.0021) | 0.0038 (0.0038) | 0.0036 (0.0034) | 0.0033 (0.0036) |
| 1_6 | 0.0035 (0.0021) | 0.0029 (0.0024) | 0.0031 (0.0022) | 0.0027 (0.0027) | 0.0029 (0.0034) |
| 1_7 | 0.0031 (0.0018) | 0.0033 (0.0027) | 0.0023 (0.0016) | 0.0030 (0.0027) | 0.0033 (0.0033) |
| 1_8 | 0.0029 (0.0019) | 0.0034 (0.0028) | 0.0030 (0.0024) | 0.0032 (0.0033) | 0.0033 (0.0036) |
| 2_1 | 0.0029 (0.0018) | 0.0031 (0.0023) | 0.0028 (0.0016) | 0.0030 (0.0036) | 0.0092 (0.0212) |
| 2_3 | 0.0027 (0.0018) | 0.0027 (0.0021) | 0.0032 (0.0026) | 0.0036 (0.0035) | 0.0030 (0.0083) |
| 2_4 | 0.0030 (0.0019) | 0.0029 (0.0025) | 0.0026 (0.0018) | 0.0030 (0.0028) | 0.0022 (0.0057) |
| 2_5 | 0.0035 (0.0020) | 0.0027 (0.0020) | 0.0027 (0.0017) | 0.0037 (0.0032) | 0.0021 (0.0029) |
| 2_6 | 0.0031 (0.0020) | 0.0030 (0.0025) | 0.0024 (0.0019) | 0.0032 (0.0030) | 0.0023 (0.0032) |
| 2_7 | 0.0030 (0.0019) | 0.0029 (0.0025) | 0.0040 (0.0036) | 0.0031 (0.0031) | 0.0018 (0.0023) |
| 2_8 | 0.0029 (0.0017) | 0.0036 (0.0028) | 0.0034 (0.0026) | 0.0038 (0.0035) | 0.0022 (0.0024) |
| 3_1 | 0.0027 (0.0018) | 0.0027 (0.0022) | 0.0030 (0.0021) | 0.0028 (0.0026) | 0.0022 (0.0056) |
| 3_2 | 0.0028 (0.0018) | 0.0029 (0.0021) | 0.0032 (0.0023) | 0.0034 (0.0034) | 0.0009 (0.0015) |
| 3_4 | 0.0028 (0.0019) | 0.0030 (0.0024) | 0.0031 (0.0024) | 0.0037 (0.0040) | 0.0054 (0.0170) |
| 3_5 | 0.0031 (0.0020) | 0.0032 (0.0024) | 0.0033 (0.0024) | 0.0033 (0.0028) | 0.0023 (0.0028) |
| 3_6 | 0.0032 (0.0019) | 0.0028 (0.0025) | 0.0033 (0.0030) | 0.0036 (0.0031) | 0.0025 (0.0037) |
| 3_7 | 0.0031 (0.0020) | 0.0031 (0.0024) | 0.0030 (0.0018) | 0.0036 (0.0035) | 0.0023 (0.0035) |
| 3_8 | 0.0032 (0.0021) | 0.0031 (0.0024) | 0.0029 (0.0026) | 0.0037 (0.0032) | 0.0026 (0.0032) |
| 4_1 | 0.0029 (0.0017) | 0.0027 (0.0022) | 0.0032 (0.0028) | 0.0032 (0.0032) | 0.0037 (0.0103) |
| 4_2 | 0.0029 (0.0021) | 0.0034 (0.0027) | 0.0024 (0.0014) | 0.0029 (0.0027) | 0.0049 (0.0109) |
| 4_3 | 0.0027 (0.0019) | 0.0032 (0.0027) | 0.0032 (0.0024) | 0.0032 (0.0030) | 0.0076 (0.0190) |
| 4_5 | 0.0030 (0.0016) | 0.0029 (0.0023) | 0.0028 (0.0024) | 0.0034 (0.0033) | 0.0028 (0.0030) |
| 4_6 | 0.0029 (0.0017) | 0.0027 (0.0021) | 0.0034 (0.0033) | 0.0033 (0.0034) | 0.0028 (0.0031) |
| 4_7 | 0.0028 (0.0017) | 0.0032 (0.0025) | 0.0029 (0.0024) | 0.0032 (0.0033) | 0.0023 (0.0028) |
| 4_8 | 0.0031 (0.0019) | 0.0028 (0.0026) | 0.0035 (0.0026) | 0.0034 (0.0036) | 0.0032 (0.0035) |
| 5_1 | 0.0030 (0.0018) | 0.0030 (0.0023) | 0.0034 (0.0030) | 0.0028 (0.0026) | 0.0033 (0.0036) |
| 5_2 | 0.0029 (0.0017) | 0.0028 (0.0023) | 0.0026 (0.0017) | 0.0036 (0.0029) | 0.0023 (0.0029) |
| 5_3 | 0.0030 (0.0018) | 0.0029 (0.0024) | 0.0029 (0.0025) | 0.0036 (0.0037) | 0.0027 (0.0030) |
| 5_4 | 0.0029 (0.0015) | 0.0026 (0.0020) | 0.0029 (0.0025) | 0.0030 (0.0033) | 0.0028 (0.0032) |
| 5_6 | 0.0027 (0.0018) | 0.0030 (0.0023) | 0.0027 (0.0016) | 0.0033 (0.0031) | 0.0074 (0.0211) |
| 5_7 | 0.0028 (0.0018) | 0.0034 (0.0029) | 0.0027 (0.0016) | 0.0034 (0.0030) | 0.0045 (0.0112) |
| 5_8 | 0.0028 (0.0017) | 0.0026 (0.0021) | 0.0032 (0.0028) | 0.0029 (0.0024) | 0.0057 (0.0110) |
| 6_1 | 0.0032 (0.0019) | 0.0031 (0.0022) | 0.0030 (0.0023) | 0.0035 (0.0030) | 0.0029 (0.0031) |
| 6_2 | 0.0031 (0.0020) | 0.0031 (0.0025) | 0.0024 (0.0018) | 0.0028 (0.0029) | 0.0021 (0.0028) |
| 6_3 | 0.0026 (0.0018) | 0.0028 (0.0024) | 0.0038 (0.0029) | 0.0031 (0.0028) | 0.0026 (0.0032) |
| 6_4 | 0.0030 (0.0019) | 0.0029 (0.0024) | 0.0034 (0.0032) | 0.0030 (0.0024) | 0.0029 (0.0031) |
| 6_5 | 0.0033 (0.0016) | 0.0032 (0.0022) | 0.0026 (0.0019) | 0.0032 (0.0029) | 0.0088 (0.0238) |
| 6_7 | 0.0029 (0.0018) | 0.0031 (0.0025) | 0.0028 (0.0023) | 0.0037 (0.0033) | 0.0025 (0.0063) |
| 6_8 | 0.0031 (0.0020) | 0.0028 (0.0027) | 0.0028 (0.0022) | 0.0030 (0.0028) | 0.0043 (0.0102) |
| 7_1 | 0.0026 (0.0017) | 0.0034 (0.0026) | 0.0024 (0.0016) | 0.0030 (0.0028) | 0.0029 (0.0028) |
| 7_2 | 0.0029 (0.0018) | 0.0027 (0.0025) | 0.0048 (0.0043) | 0.0035 (0.0031) | 0.0018 (0.0024) |
| 7_3 | 0.0034 (0.0023) | 0.0029 (0.0024) | 0.0030 (0.0019) | 0.0034 (0.0028) | 0.0023 (0.0028) |
| 7_4 | 0.0029 (0.0017) | 0.0031 (0.0024) | 0.0028 (0.0021) | 0.0030 (0.0031) | 0.0023 (0.0030) |
| 7_5 | 0.0031 (0.0018) | 0.0033 (0.0024) | 0.0029 (0.0017) | 0.0030 (0.0029) | 0.0039 (0.0094) |
| 7_6 | 0.0030 (0.0020) | 0.0029 (0.0026) | 0.0027 (0.0022) | 0.0037 (0.0035) | 0.0030 (0.0068) |
| 7_8 | 0.0028 (0.0020) | 0.0025 (0.0021) | 0.0025 (0.0018) | 0.0033 (0.0029) | 0.0059 (0.0182) |
| 8_1 | 0.0029 (0.0019) | 0.0030 (0.0026) | 0.0034 (0.0026) | 0.0037 (0.0035) | 0.0032 (0.0037) |
| 8_2 | 0.0027 (0.0018) | 0.0027 (0.0021) | 0.0031 (0.0025) | 0.0036 (0.0029) | 0.0023 (0.0030) |
| 8_3 | 0.0030 (0.0018) | 0.0035 (0.0027) | 0.0032 (0.0024) | 0.0034 (0.0033) | 0.0025 (0.0027) |
| 8_4 | 0.0028 (0.0019) | 0.0027 (0.0023) | 0.0031 (0.0025) | 0.0041 (0.0042) | 0.0031 (0.0035) |
| 8_5 | 0.0031 (0.0022) | 0.0032 (0.0027) | 0.0033 (0.0028) | 0.0036 (0.0035) | 0.0047 (0.0111) |
| 8_6 | 0.0030 (0.0021) | 0.0032 (0.0027) | 0.0028 (0.0024) | 0.0035 (0.0032) | 0.0043 (0.0117) |
| 8_7 | 0.0030 (0.0020) | 0.0030 (0.0024) | 0.0024 (0.0015) | 0.0033 (0.0038) | 0.0063 (0.0180) |

## Supplementary Figures

**Figure S1.**
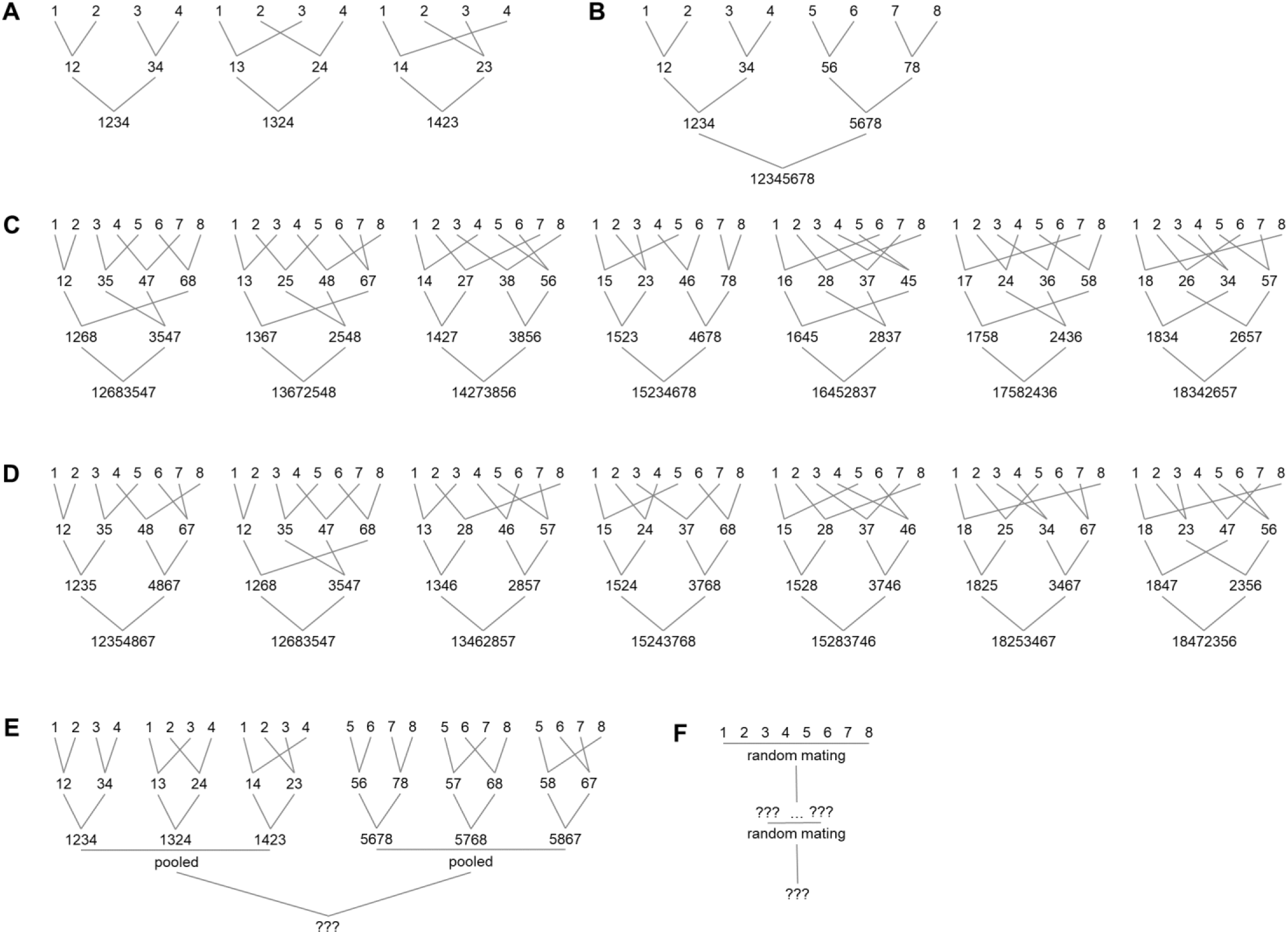
Examples of MAGIC population designs. [**A**] Full design with 4 founders. [**B**] Basic design with 8 founders. [**C**] Partial balanced design with 7 funnels and 8 founders. [**D**] Partial unbalanced design with 7 funnels and 8 founders. [**E**] Semi-structured design with 8 founders. [**F**] Unstructured design with 8 founders.

**Figure S2.**
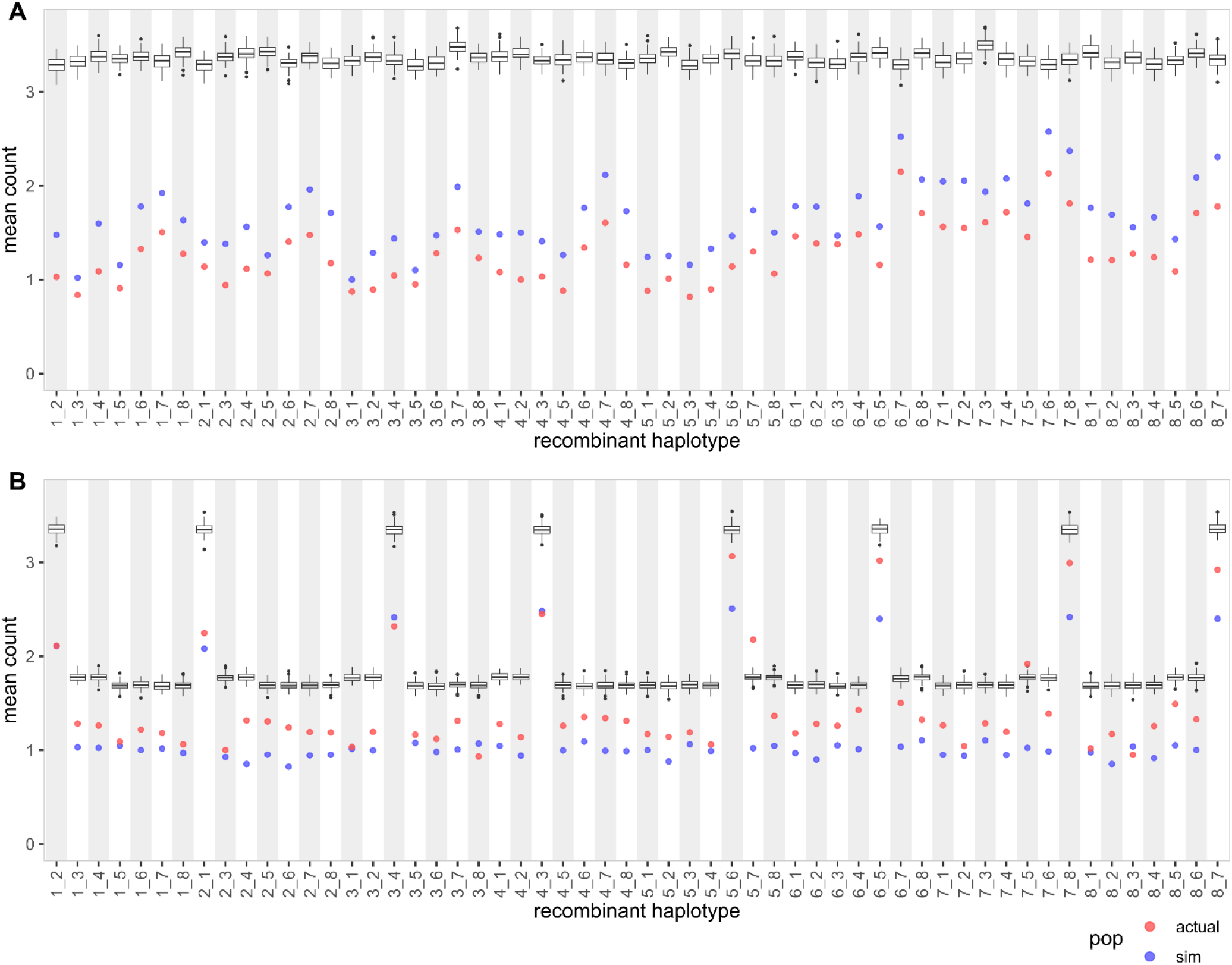
Distributions of recombinant haplotypes in two wheat MAGIC populations (full datasets). [**A**] Plot shows mean count of each recombinant haplotype in a single RIL in wheat-UK8 (full dataset with 643 RILs and 18,599 markers). The boxplot shows mean count from true founder genotypes (100 simulated iterations). The red and blue points show mean count from inferred founder genotypes. [**B**] Plot shows mean count of each recombinant haplotype in a single RIL in wheat-DE8 (full dataset with 910 RILs and 7,579 markers).

**Figure S3.**
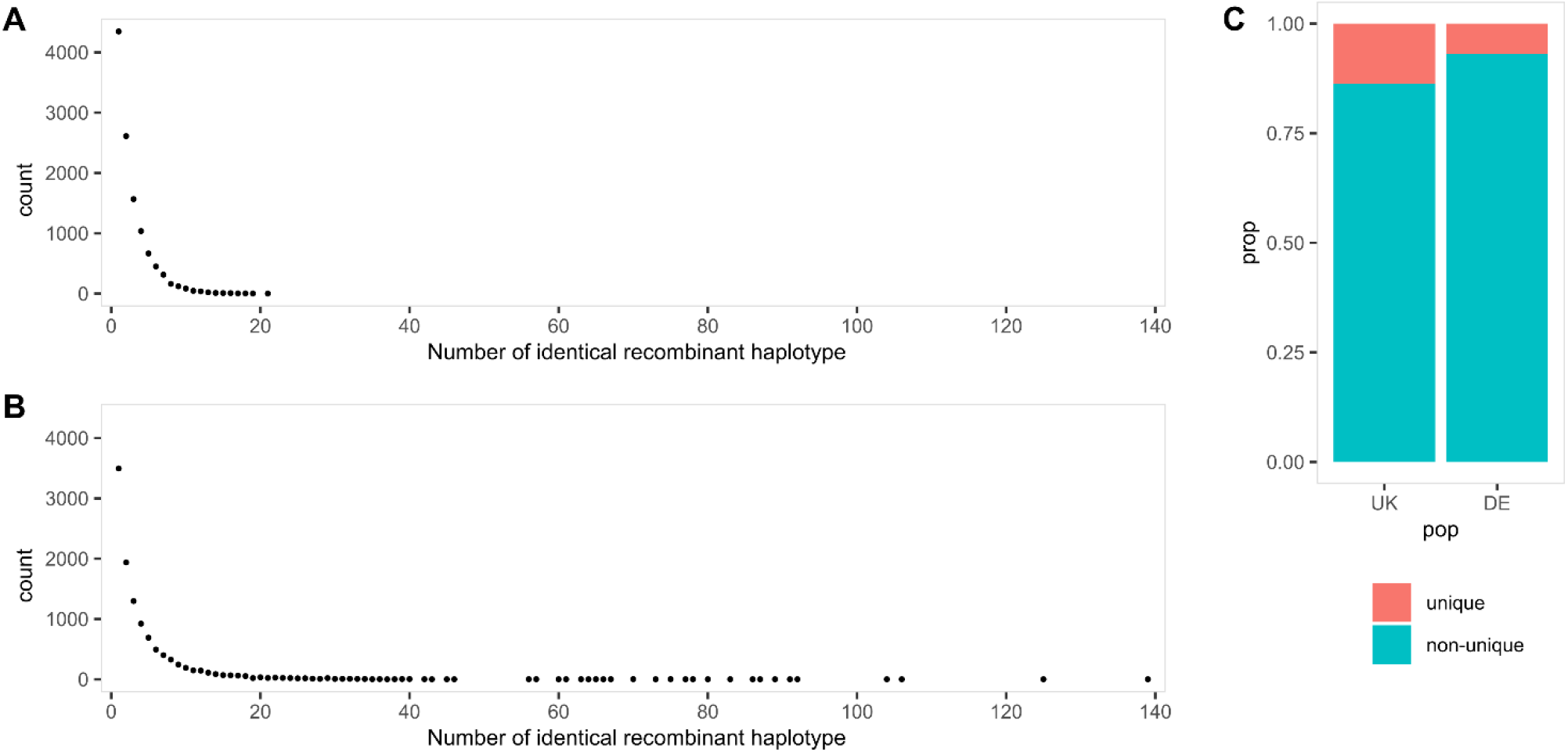
Distributions of unique and identical recombinant haplotypes in two wheat MAGIC populations. Recombinant haplotypes are considered identical if they are of the same founder pairs and present in the same 10 cM interval, otherwise unique. [**A**] Counts of the number of identical recombinant haplotypes in wheat-UK8. The left most point is the count of unique recombinant haplotypes. [**B**] Counts of the number of identical recombinant haplotypes in wheat-DE8. [**C**] Proportions of unique and non-unique (identical) recombinant haplotypes in wheat-UK8 and wheat-DE8.

**Figure S4.**
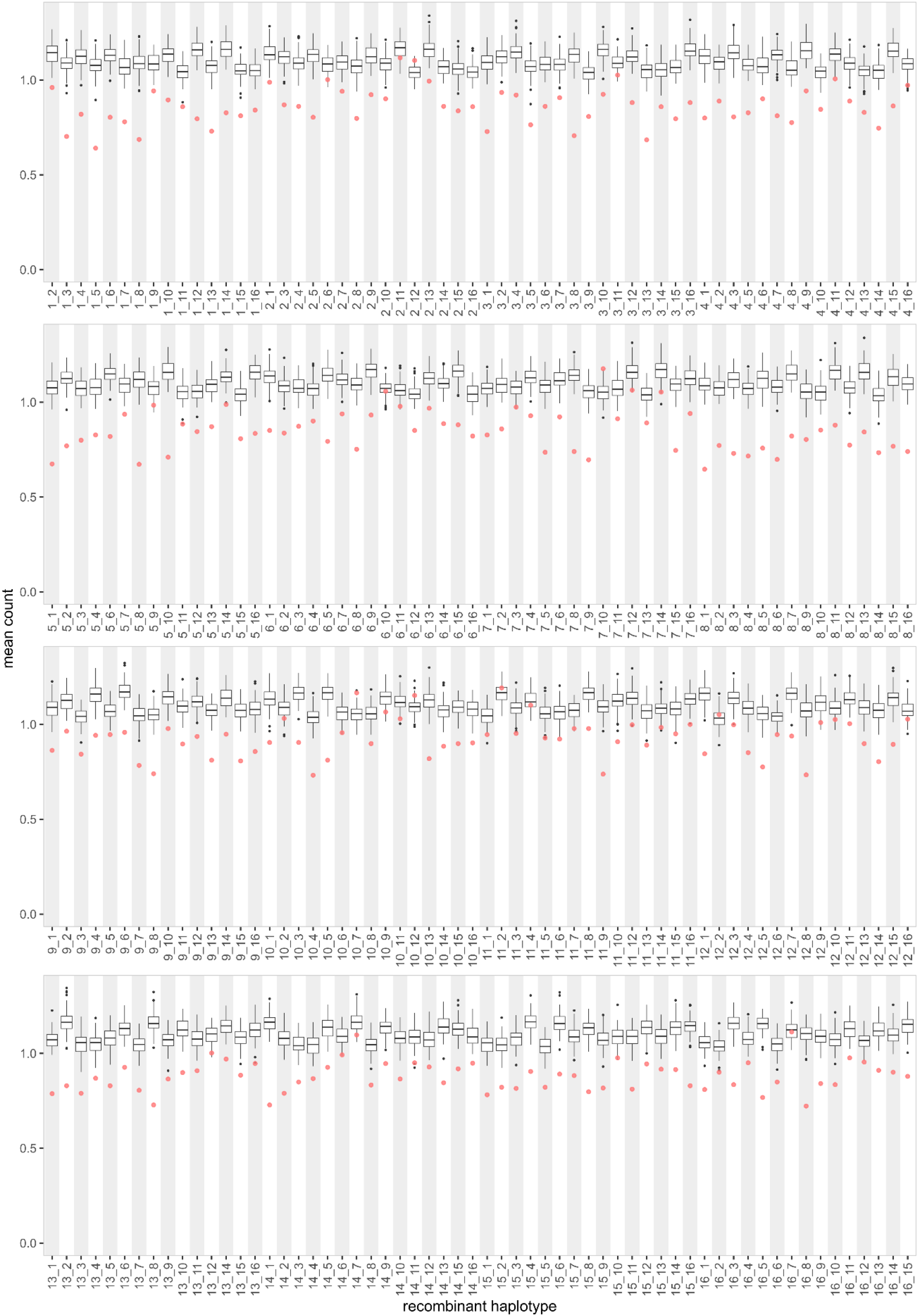
Distributions of recombinant haplotypes in wheat-UK16. Plot shows mean count of each recombinant haplotype in a single RIL in wheat-UK16. The boxplot shows mean count from true founder genotypes (100 simulated iterations). The red points show mean count from inferred founder genotypes in actual dataset.

